# Rad51, Rad54, and ZMM proteins antagonize the mismatch repair system to promote fertility of budding yeast intraspecies hybrid zygotes

**DOI:** 10.1101/2025.02.08.636946

**Authors:** Ya-Ling Hung, Chi-Ning Chuang, Hong-Xiang Kim, Hou-Cheng Liu, Jhong-Syuan Yao, Lavernchy Jovanska, Yi-Ping Hsueh, Ruey-Shyang Chen, Ting-Fang Wang

## Abstract

Rad51 and meiosis-specific Dmc1 catalyze homologous recombination (HR) between maternal and paternal chromosomes during meiosis in many sexual eukaryotes, generating three interhomolog (IH) recombination products: noncrossovers (NCOs), class I crossovers (COs), and class II COs. Some COs form chiasmata, which physically connect homologous chromosomes and ensure proper chromosome segregation during meiosis I. Meiosis is highly relevant to speciation, with the mismatch repair (MMR) system believed to prevent IH-HR, leading to postzygotic isolation between closely related species. We report that several *Saccharomyces cerevisiae* HR proteins exhibit anti-MMR activities, including Rad51, Rad54, and synapsis-promoting ZMM proteins (Mer3, Zip1, Zip4, and Msh4) in SK1/S288c hybrid zygotes. Srs2 (an ortholog of *Escherichia coli* helicase UvrD) facilitates MMR by dissembling Rad51-ssDNA presynaptic filaments. Rad51 antagonizes MMR and Srs2 to catalyze D-loop formation. Rad54’s anti-MMR activity acts after Srs2 and outcompetes its pro-HR function to promote Rad51-mediated IH-HR in hybrid zygotes. Dmc1 does not possess anti-MMR activity, but exhibits better mismatch tolerability than Rad51. Following D-loop formation mediated by Dmc1 and/or Rad51, ZMM proteins promote class I IH-CO formation while limiting MMR to promote NCO formation by Sgs1 (an ortholog of *E. coli* RecQ helicase) and prevent class II IH-CO formation by the Mms4•Mus81 endonuclease.

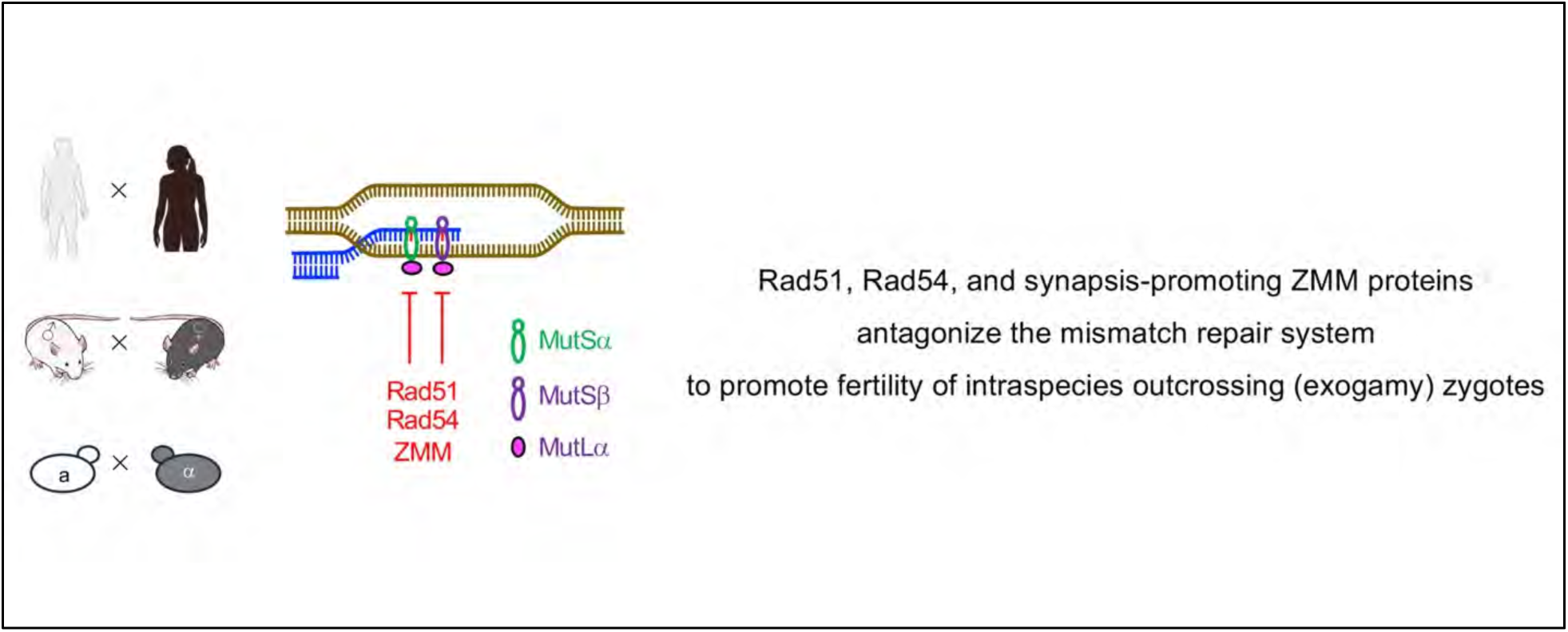

**Highlights:**

- MMR and HR antagonize each other in intraspecies hybrid zygotes.
- The HR proteins Rad51, Rad54, and ZMM exhibit anti-MMR activities.
- MMR overrides HR in hybrid zygotes with anti-MMR mutations, causing hybrid sterility.

## Introduction

Meiosis, a specialized cell division of sexual eukaryotes, generates haploid gametes from diploid precursor cells. This process occurs via one round of DNA replication and two sequential nuclear divisions, i.e., meiosis I (MI) and meiosis II (MII). After replication, the parental and maternal chromosomes each produce two sister chromatids, which are paired and held together via cohesion complexes. The sister chromatid pairs then interact and form a bivalent that encompasses a total of four copies of double-stranded DNA (dsDNA). This process requires the formation of chiasmata via interhomolog (IH) homologous recombination (HR) to physically connect homologous chromosomes and ensure their correct segregation during MI.

In most studied sexual eukaryotes, HR in meiosis begins between premeiotic S phase and meiotic prophase I with the introduction of double-strand breaks (DSBs) by the highly conserved endonuclease Spo11 and its accessory proteins (1). The bound Spo11 enzyme blocks the nonhomologous end joining (NHEJ) DSB repair pathway by preventing binding of the heterodimeric Ku70•Ku80 complex (2,3). In the budding yeast *Saccharomyces cerevisiae* (the subject of this study), DSBs are first resected 5′ to 3′ to form long single-stranded DNA (ssDNA) overhangs, with a median length of gene conversion close to 2000 base pairs (bps) (4) or 1000 bps (5). These ssDNAs are then protected by replication protein A (RPA). Two RecA-like recombinases (ubiquitous Rad51 and meiosis-specific Dmc1) cooperate to repair DSBs preferentially via the IH-HR pathway rather than the intersister (IS)-HR pathway, i.e., using a homologous non-sister chromosome rather than a sister chromatid as a template. This process involves invasion of an intact template and formation of a displacement loop (D-loop), followed by DNA repair synthesis of the free 3’ end in the D-loop. Rad51 and Dmc1 do not act alone. Rad51 has several accessory factors, such as the PCSS complex (Psy3•Csm2•Shu1•Shu2), Rad52, Rad54, the Rad55•Rad57 heterodimer, Rad59, Srs2, Pif1 and Hed1. Dmc1’s accessory factors include Rad51, Tid1 (also called Rdh54), the Mei5•Sae3 heterodimer, and the Hop2•Mnd1 heterodimer (6). The Psy3•Csm2 complex is structurally similar to the Rad51 dimer and can stabilize the Rad51-ssDNA nucleoprotein filaments (7). Rad52 exerts dual roles. First, it associates with RPA and functions with the Rad55•Rad57 heterocomplex to assemble Rad51-ssDNA nucleoprotein filaments (8). Second, the single strand annealing (SSA) activity of Rad52 promotes second-end capture in DSB repair to form stabilized joint molecules, ultimately leading to the generation of Holliday junctions (9,10). Rad59 interacts with Rad52 to regulate associations of Rad52 with DSBs (11) and also to augment Rad52’s activity in the SSA pathway (12). Srs2 is a member of the superfamily 1 (Sf1a) helicases that move in the 3′ → 5′ direction relative to the bound strand of nucleic acid (13,14). Srs2 binds to Rad51 (not Dmc1), competitively ejects Rad52 and the Rad55•Rad57 complex, and then disassembles the Rad51-ssDNA presynaptic filament (15–18). Unlike Rad52, the PCSS complex does not appear to promote Rad51 function during meiosis by antagonizing Srs2 (18). The Rad51 patches on ssDNA can stimulate filament assembly of Dmc1 *in vitro* (19), with Dmc1 being a potent inhibitor of Srs2 (20). *S. cerevisiae* Rad54 and Tid1 are members of the Swi2/Snf2 family of dsDNA-dependent ATPases that specifically facilitate the homology pairing and strand exchange activities of their biological partners Rad51 and Dmc1, respectively (21). Rad54 is a multifunctional actor in Rad51-mediated HR. First, it binds tightly to Rad51-associated ssDNA and acts downstream of Srs2 to promote Rad51-mediated D-loop formation (22). Second, it exerts chromatin remodeling activity to clear nucleosomes or other proteins during homology pairing (23). Third, Rad54 also possesses Rad51-dependent branch migration activity to dissociate Rad51-generated D-loops (24). Dmc1-made D-loops are more resistant to Rad54’s branch migration activity than those formed by Rad51 (25). Tid1 translocates on DNA with a similar processivity (∼10,000 bp) and complexity as Rad54, but at ∼2–4-fold lower speed, so it has been proposed that Tid1 may disrupt Dmc1-made D-loops (26). The Rad51-A320V, Rad51-Y388H, and Rad51-G393D mutant proteins are defective in binding to Rad52 and/or Srs2, yet they retain the ability (equivalent to wild-type) to interact with Rad54, Tid1, and Rad59 (27). Hed1 is a meiosis-specific inhibitor of Rad51 (28) that prevents Rad51 from interacting with Rad54 (29). Mph1 is the *S. cerevisiae* ortholog of the mammalian Fanconi anemia complementation group M (FANCM) helicase that works independently of Srs2 to dissociate Rad51-made D-loops during vegetative growth (30). Mph1 also prevents precocious DSB strand exchange between sister chromatids before homologs have completed pairing, thereby ensuring high levels of IH-HR products between non-sister homologous chromosomes in meiosis (31).

It has been nicely demonstrated that *S. cerevisiae* meiotic recombination occurs in two temporally distinct phases (32). During Phase 1, Rad51 is inhibited by Hed1, and Dmc1 mediates IH-HR and promotes homolog synapsis, i.e., assembly of the synaptonemal complex (SC), an evolutionarily conserved protein lattice that connects paired homologous chromosomes in meiotic prophase (33,34). In Phase 2, Rad51 becomes active and functions with Rad54 to repair residual DSBs, making increasing use of sister chromatids. Pif1, an ATP-dependent 5′-to-3′ DNA helicase, specifically promotes Rad51-mediated IH-HR, but not Dmc1-mediated IH-HR (32). The transition from Phase 1 to Phase 2 is controlled by a meiotic checkpoint through the action of a meiosis-specific effector kinase Mek1 and two meiosis-specific chromosomal proteins (Red1 and Hop1) to uphold the IH bias of meiotic recombination (35–41). Activation of Mek1 requires DSB- and Red1-dependent Hop1 phosphorylation at threonine 318 (T318) (40,42,43). Hop1-T318 phosphorylation is mediated by Tel1^ATM^ and Mec1^ATR^, the yeast homologs of the mammalian DNA damage sensor kinases ATM (ataxia-telangiectasia mutated) and ATR (RAD3-related) (44). Mek1 downregulates Rad51 activity in the *dmc1*Δ mutant line by phosphorylation of Hed1 at residue T40 (45). Mek1 also phosphorylates Rad54 at residue T132 to regulate recombination partner choice in meiosis. First, it reduces Rad51 activity by inhibiting Rad51•Rad54 complex formation and, second, it suppresses Rad51-mediated strand invasion of sister chromatids via a Rad54-independent mechanism (46). When only Rad51 is present in meiotic cells (e.g., *dmc1*Δ (47) and *dmc1*Δ *hed1*Δ (48) cells), most DSBs are not properly repaired during Phase I, leading to chronic overactivation of the DNA damage checkpoint pathway and persistent Hop1-T318 phosphorylation (48).

ATR^Mec1^ and ATM^Tel1^ also phosphorylate the NH_2_-terminal domain (NTD; residues 1–66) of *S. cerevisiae* Rad51 during vegetative growth and meiosis. ATR^Mec1^/ATM^Tel1^-dependent NTD phosphorylation antagonizes Rad51 degradation via the proteasomal pathway, increasing the half-life of Rad51 from **∼**30 min to ≥180 min. Rad51-ΔN is a hypomorphic mutant lacking the NTD. The three serine residues of the serine(S)-glutamine (Q) motifs (S2Q, S12Q, and S30Q) of Rad51-NTD3A are mutated into alanines (As). Steady-state Rad51 protein levels in meiotic *rad51-*ΔN and *rad51-NTD3A* cells are only ∼3% and ∼30% that of Rad51 in wild-type (WT) cells, explaining why Mec1^ATR^ plays an important role in regulating protein homeostasis in *S. cerevisiae* (49). Rad51-ΔN and Rad51-NTD3A proteins, like wild-type Rad51, remain catalytically active in homology-directed DSB repair during meiosis because *rad51*Δ, *rad51-*Δ*N*, and *rad51-NTD3A* meiotic cells produce <1%, ∼50% and ∼97% viable spores, respectively (48).

The IH-HR pathway ultimately produces two recombination products, i.e., crossovers (COs) and non-crossovers (NCOs; also called gene or allelic conversion). IH-COs are the primary source of genetic diversity in gametes, whereas IH-NCOs involve only the unidirectional transfer of genetic information. Some IH-COs result in chiasmata and, together with sister chromatid cohesion, allow spindle tension to be generated when homologous chromosomes begin to segregate during MI (see review in (6)). Homology-directed DSB repair in both NCOs and COs involves DNA repair synthesis, often with multiple rounds of strand invasion (50). In *S. cerevisiae*, IH-NCOs form earlier than IH-COs and are mainly produced via the synthesis-dependent strand annealing (SDSA) pathway (51–53). IH-NCO formation is abolished during early meiosis in *S. cerevisiae* mutants lacking Sgs1 (the yeast homolog of *E. coli* and mammalian RecQ helicases) (54). It is not entirely clear how Sgs1 is recruited to DSBs during early meiosis and promotes IH-NCO formation. CO-specific double Holliday junction formation occurs via processes (e.g., DNA repair synthesis and repeated strand exchange) involving branch migration as an integral feature, which can be separated from repair of the break itself (31). If homologs harbor different alleles, the resulting heteroduplex DNA (hDNA) by IH-HR will contain mismatched base pairs. The mismatched base pairs in hDNA are detected genetically as departures from Mendelian segregation (4^+^:4^−^, based on the eight ssDNA strands present) in haploid meiotic products. When allelic gene conversion occurs during meiosis between heterozygotic sites, mismatches on only one homolog (asymmetric heteroduplex) produce 5^+^:3^−^ or 3^+^:5^−^ marker segregation (half conversion), as well as 6^+^:2^−^ or 2^+^:6^−^ marker segregation (full conversion). The mismatched base pairs in hDNA are then recognized and/or corrected by the mismatch repair (MMR) system (see below), thus causing one of the alleles to be converted to the other. This process can cause non-Mendelian segregation (NMS), i.e., 3^+^:1^−^, 1^+^:3^−^, 4^+^:0^−^, or 0^+^:4^−^ alleles in tetrads (based on the four dsDNA duplexes present). In contrast, Mendelian segregation (MS) results in 2^+^:2^−^ tetrads (50,51,55–57).

Most proteins involved in the prokaryotic and eukaryotic MMR systems are conserved both structurally and functionally. During the MMR process in *E. coli*, mismatched base pairs in hDNA are first recognized by the MutS homodimer. Subsequently, a ternary complex comprising DNA, MutS, and MutL is formed in a reaction requiring ATP. A nick is then introduced by MutH into the unmethylated DNA strand at a hemimethylated d(GATC) site. Next, the ATP-bound MutL dimer loads the UrvD helicase onto the nicked DNA. The UvrD loading event ensures that the helicase is loaded onto the correct strand to translocate in the 3′ to 5′ direction toward the mismatched based pair (58). Mismatch recognition in most studied eukaryotes is trigged by two evolutionarily conserved MutS complexes, i.e., Msh2•Msh3 (MutSα) and Msh2•Msh6 (MutSα). MutSα is primarily involved in repairing base-base and small insertion/deletion loop mismatches, whereas MutSα acts primarily on insertion/deletion loop mismatches up to 12 nucleotides in length (59). During mismatch correction, the MutLα (Mlh1•Pms1) endonuclease is then activated in a mismatch-dependent manner by MutSβ (or MutSα), RPA, PCNA, RFC, and DNA polymerase δ in reconstituted 3′-nick-directed MMR reactions (60). MutLα functions as the primary factor in repairing mismatched base pairs (61). MutLα (Mlh1•Mlh2) is a non-essential accessory factor that acts to enhance the activity of MutLα during vegetative growth (62). Using a yeast strain with an HO endonuclease-induced DSB that can be repaired by single-strand annealing (SSA) between flanking homologous sequences, heteroduplex rejection was found to be mediated by Sgs1 and Msh2, but not by the yeast MutL proteins. Moreover, correction of mismatched base pairs within the SSA intermediates required MutLα to the same extent as Msh2 (63). During meiosis, MutLβ forms a dynamic ensemble with a meiosis-specific DNA helicase Mer3 at the Dmc1-made D-loops to limit Sgs1-dependent NCO *in vivo* (64,65). MutLγ is a Msh2-stimulated endonuclease that mediates CO-specific resolution of double Holliday junctions (66–71). UvrD can dismantle RecA nucleoprotein filaments in *E. coli* (72). The primary function of Srs2, the yeast paralog of UvrD, is to dissemble the Rad51-ssDNA presynaptic filament during HR (73). It remains unclear if Srs2 is also involved in yeast MMR or nucleotide excision repair.

Two mechanisms have been reported to regulate chromosome-wide IH-CO positions (74–77). First, the “CO assurance rule” reflects the fact that each pair of homologous chromosomes must receive at least one obligatory chiasma during the diplotene stage of meiosis prophase I because a chiasma ensures proper chromosome segregation at MI. Second, IH-COs too close to each other might result in diminished sister chromatin cohesion and tension at MI. The “CO interference rule” reflects the fact that superfluous IH-COs lead to chromosome nondisjunction (NDJ) and aneuploidy. In classical genetics, interference (γ) is defined as 1 − (coefficient of coincidence; c.o.c.), where c.o.c is calculated by dividing the actual double recombinant frequency by the expected double recombinant frequency. Within many sexual eukaryotes, two subclasses of IH-COs co-exist. The class I IH-CO pathway accounts for 70-85% of COs in *S. cerevisiae* (51) and requires a group of meiosis-specific ZMM proteins, including Zip1, the Zip2•Zip4•Spo16 (ZZS) complex, Mer3 DNA helicase, the Msh4•Msh5 (MutSγ) complex, MutLγ, and Zip3 (51,64,66–69,71,78–84). This group of IH-CO products is interference-sensitive or interfering, with formation of COs in relative proximity to each other appearing to be suppressed (51,85). The ZMM proteins are known to antagonize Sgs1’s potent meiotic anti-CO activity (86) and they are also termed the synaptic initiation complex (SIC) because they control homolog synapsis or SC assembly (79). Zip1 is the main structural component of the SC central element (87) and it antagonizes Mph1-mediated dissociation of late-occurring joint molecules (31). The ZZS complex performs multiple distinct activities. Zip4 acts as a hub for physical interactions with components of the chromosome axis (e.g., Red1 and Hop1), with components of the SC central element (e.g., the Ecm11•Gmc2 complex), and with the crossover machinery (88,89). The Zip4-N919Q mutant protein specifically abolishes the Zip4-Ecm11 interaction, leading to loss of Ecm11 binding to chromosomes and defective SC assembly (89). Spo16 binds the C-terminal XPF domain of Zip2 to form a meiosis-specific complex that shares structural similarities to the structure-specific XPF–ERCC1 nuclease, although it lacks endonuclease activity. The Zip2-ΔXPF mutant protein lacks the XPF domain, which mediates the non-catalytic DNA recognition function of the Zip2•Spo16 complex that promotes IH-CO formation via binding to joint molecule recombination intermediates. Despite a minor difference in CO levels, both sporulation efficiency and spore viability of the *zip2-*Δ*XPF* mutant are significantly greater than those of the *zip2*Δ mutant (90,91). *S. cerevisiae* Zip3, and its paralogs RNF212 and HEI10 in other sexual eukaryotes, are E3 ligases of ubiquitin and/or small ubiquitin-like modifier (SUMO) (51,80–84). Zip3 and MutLγ are enriched at class I IH-COs, appearing as bright foci on pachytene chromosomes (76,80,83,84,92). Mer3 is a meiosis-specific DNA helicase displaying several functions. Mer3 physically interacts with Dmc1 to antagonize the anti-recombinase activities of Sgs1 (65), and thus selectively affects IH-CO but not IH-NCO formation. Mer3 also interacts with MutLα to compete with the Pif1 5′-to-3′ DNA helicase for binding to the PCNA clamp loader RFC, thereby limiting polymerase delta (Pol δ)-mediated DNA synthesis during DSB repair synthesis in meiosis (93). Pif1 specifically promotes Rad51-mediated IH-HR but not Dmc1-mediated IH-HR (32). Mer3 can stimulate 3′-to-5′ heteroduplex extension of Rad51, thereby stabilizing nascent joint molecules via DNA heteroduplex extension to permit capture of the second processed end of a dsDNA break, a step required for the formation of IH-CO products (52). Although *dmc1*Δ *hed1*Δ results in unrepaired DSBs and persistent hyperactivation of the meiotic checkpoint (48), the Rad51-Mer3 interaction may explain why SC formation in *dmc1*Δ *hed1*Δ meiotic cells is inefficient and abnormal but not completely blocked (94). Msh4 and Msh5 form a meiosis-specific MutSγ heterodimer that is not involved in MMR or recognizing mismatched base pairs, but it does facilitate crossing over by binding and stabilizing nascent recombination intermediates (91,95,96). It has also been reported that CO assurance requires all ZMM proteins and a full-length SC. In contrast, CO interference requires the normal assembly of recombination complexes containing MutSγ, but it does not require ZZS- or Zip3-dependent extension of the SC (97).

Class II IH-COs were classically believed to be non-interfering in several sexual eukaryotes (75,85,98). However, a recent study has indicated that they might be interfering in *S. cerevisiae* (99). This pathway is independent of ZMM proteins and MutLγ, but involves the Mms4•Mus81 endonuclease complex (85) and, to a lesser degree, Yen1/GEN1 resolvase (Holliday junction 5’ flap endonuclease) and the Slx1•Slx4 endonuclease heterocomplex (100,101). Srs2 was reported previously to promote Mms4-mediated resolution of D-loops (102–105), yet it has been shown using a Srs2 separation-of-function mutant that the ability of Srs2 to remove Rad51 from ssDNA is its primary role during HR (73). It was also reported that Mms4•Mus81 might collaborate with Sgs1 to promote joint molecule resolution *in vivo* and to direct meiotic recombination toward IH interactions that promote proper chromosome segregation (106,107). The *sgs1*Δ-C795 *S. cerevisiae* cell line, in which the carboxy-terminal 795 amino acids of Sgs1 have been removed, represents a separation-of-function mutant that defines the function of Sgs1 that is important for *S. cerevisiae* meiosis (108). In sexual eukaryotes (e.g., the fission yeast *Schizosaccharomyces pombe* and the free-living ciliate *Tetrahymena thermophila*) that have lost the SC during their evolution, Sgs1 and Mms4•Mus81 are sufficient to execute meiotic recombination (106,107,109). Not only does Sgs1 prevent aberrant crossing-over by suppressing formation of multichromatid joint molecules, but it also functions as an IH-CO-promoting factor (50,54,68,86,108,110–112).

Meiosis is highly related to speciation. Hybrid zygotes between closely related species are often sterile or inviable. Hybrid infertility is an important feature of the speciation process, acting as a reproductive isolating barrier. For almost 30 years, it was believed that the two haploid genome sequences in hybrid zygotes are too diverged to cross over efficiently during meiosis, resulting in MI nondisjunction (NDJ) and aneuploidy. The MMR system was thought to be the major impediment to IH-HR during meiosis in sexual eukaryotes and interspecific sexual conjugation in prokaryotes, as the requirement for DNA sequence homology during sexual reproduction is relaxed in mutants deficient in anti-recombination genes (57,113–127). The mismatched based pairs in hDNA are recognized and corrected by the MMR system via removal and resynthesis of strands over large regions. Mismatch recognition triggers heteroduplex correction and/or rejection, whereby helicases disassemble recombination intermediates that contain heteroduplexes. Deleting *MSH2* inactivates both heteroduplex rejection and MMR, thereby preserving heteroduplex patterns and allowing inference of parental strand contributions from genotypes of spores in tetrads (4,113,114,116–121). Sgs1 acts downstream of Msh2-mediated mismatch recognition to unwind nascent recombination intermediates containing a high density of heteroduplex base pair mismatches (127). Accordingly, the fertility of interspecies *Saccharomyces cerevisiae* W303/*Saccharomyces paradoxus* N17 hybrid zygotes can be increased to the level of non-hybrids by introducing *P_CLB2_-SGS1* (86,107,128,129) and *P_CLB2_-MSH2 P_CLB2_-SGS1* (129). The genomic sequences of *S. cerevisiae* and *S. paradoxus* differ by >12% (129). *P_CLB2_* is the native promoter of the mitosis-specific B-type cyclin gene *CLB2* that is deactivated during meiosis (130), thereby specifically repressing *MSH2* and *SGS1* during that cell division (86,107,128,129). Since W303/N17 hybrid zygotes possessing *P_CLB2_-SGS1* and *P_CLB2_-MSH2 P_CLB2_-SGS1* can effectively produce viable spores, hybrid sterility and allelic incompatibility are unlikely to occur in these zygotes, with unexpected consequent effects for important protein-protein interactions (131).

Most of our current knowledge about *S. cerevisiae* meiotic recombination was mainly derived from studies of purebred diploid strains (e.g., SK1/SK1, BR/BR or S288c/S288c) and intraspecies hybrid diploid strains (e.g., SK1/S288c, SK1/BR or S288c/BR) because they are fertile and produce many viable spores (4,127). The nucleotide sequences (or genotypes) of two haploid genomes in purebred zygotes are nearly identical to each other due to long-term inbreeding, whereas those in intraspecies hybrid zygotes often possess a significant degree of sequence heterogeneity. For example, the near-complete genomic sequences of *S. cerevisiae* SK1 and S288c haploid strains harbor 225,265 bps (1.85%) of single nucleotide polymorphisms (SNPs) and 988,818 bps (8.14%) of insertions and deletions (InDels) (5,132,133). In Supplemental Datasets 1-17 (DS1-DS17) of the current report, we provide all of the SNPs and InDels between the 16 chromosomes and near-complete genomes of SK1(*MATa*) and S288c(*MATα*) haploid cells. Like W303/N17 hybrid zygotes, SK1/S288c hybrid zygotes are unlikely to exhibit allelic incompatibility due to their high fertility, which could have unexpected effects on important protein-protein interactions (131). Genome-wide mapping of meiotic recombination products (in tetrads with four viable spores) revealed that deletion of *MSH2* from SK1/S288c hybrid zygotes only increased COs by approximately 20% (4).

In this report, we propose that HR and MMR are yin-yang balanced partners during meiosis of SK1/S288c intraspecies hybrid zygotes, and that some HR proteins can antagonize the MMR system in those zygotes. This yin-yang relationship is biased toward IH-HR in purebred zygotes (e.g., SK1/SK1), but toward MMR in the anti-MMR mutants of SK1/S288c hybrid zygotes. Accordingly, HR proteins with anti-MMR activities exhibit two distinct properties. First, they are preferentially required for fertility (i.e., the formation of viable spores) of SK1/S288c hybrid zygotes compared to SK1/SK1 purebred zygotes. Second, the low fertility (spore viability) phenotype of the anti-MMR mutants of SK1/S288c hybrid zygotes can be rescued (at least partly) by removal or meiotic repression of MMR genes (e.g., *MSH2*). We have discovered several anti-MMR mutants of IH-HR genes that regulate the yin-yang relationship between HR and MMR in SK1/S288c hybrid meiosis. The anti-MMR activities of HR proteins display two roles: first, they prevent the MMR system from recognizing mismatched base pairs during D-loop formation; and second, they remove the MMR system from mismatched base pairs in D-loops and the resulting joint molecules.

## Materials and Methods

### Yeast strains and cultures

Standard methods were deployed for the growth and genetic manipulation of yeast (134). Unless noted otherwise, all *S. cerevisiae* (5,48) strains were grown at 30 °C in YPD medium (1% yeast extract, 2% peptone, and 2% glucose). KanMX, HphMX, and NatMX cassettes, which confer resistance to G418, hygromycin or nourseothricin, respectively (135), were incorporated into the SK1/SK1 purebred and SK1/S288c hybrid strains. G418, hygromycin, and nourseothricin were added to the media at final concentrations of 200 μg/ml, 300 μg/ml, and 50 μg/ml, respectively (136). All meiotic experiments were performed as described previously (82,137) using SK1/SK1 purebred diploid cells and SK1/S288c hybrid diploid cells (DS18). Figure 1A shows a schematic diagram of specific DSB hotspots in three haploid strains, respectively. Both SK1(*a, VIII, rgC*)/SK1(*α, VIII, RGc*) purebred zygotes and S288c(*a, VIII, rgC*)/SK1(*α, VIII, RGc*) hybrid zygotes carry three heterozygous spore-autonomous fluorescent marker genes (*RFP, GFP*, and *CFP*) (DS18) (138), with *MATa, MATα*, chromosome VIII, *CEN8::RFP, CEN8*, *ARG4::GFP*, *ARG4, THR1::CFP* and *THR1* indicated as *a*, *α*, *VIII, R*, *r*, *G*, *g, C*, and *c,* respectively.

**Figure 1.**
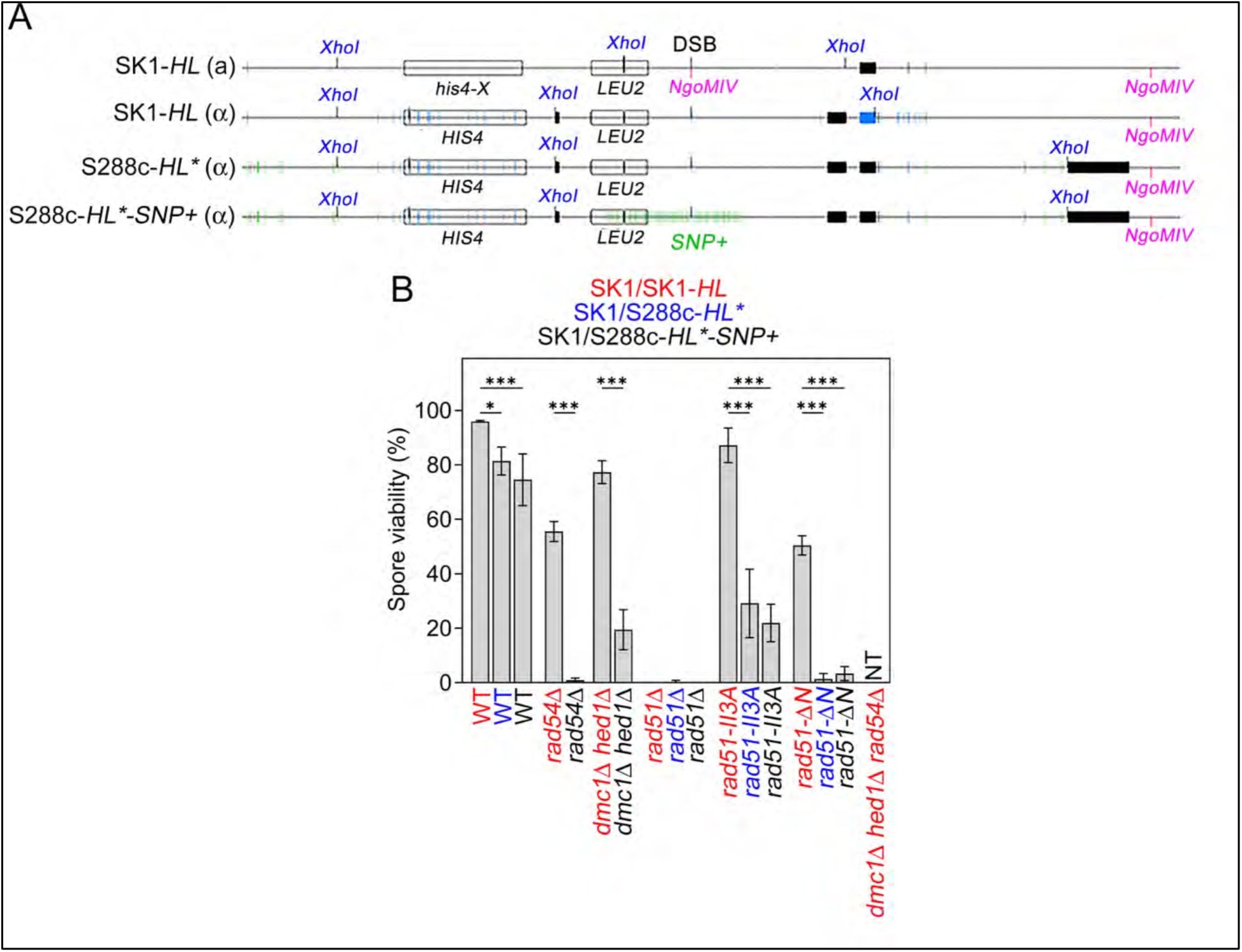
Validation of intraspecies purebred and hybrid strains used for screening anti-MMR mutants. **(A)** Schematic of the *HIS4-LEU2* DSB hotspots in SK1/SK1-*HL* purebred zygotes, and SK1/S288c-*HL*\* and SK1/S288c-*HL*\* *-SNP+* hybrid zygotes. The α cell-specific and S288c-specific SNPs are indicated in blue and green, respectively. Deletions are indicated in black. **(B)** Spore viability. SK1/SK1-*HL* purebred zygotes (in red), SK1/S288c-*HL*\* hybrid zygotes (in blue) and SK1/S288c-*HL*\* *-SNP+* hybrid zygotes (in black) are indicated in red, blue, and black, respectively. Yeast cells were grown in YPDA medium and then transferred into sporulation medium at 30 °C for 48 hours. The tetrads were subjected to limited zymolyase digestion, sufficient to break down the protective membrane surrounding spores. The digested tetrads were transferred onto the surface of an YPDA agar plate on a microscopic platform. Tetrads were selected, picked up, and their individual spores were pulled apart using a micromanipulator device controlling the movement of a fine glass needle. Both hybrid zygotes and the purebred zygotes produced 77.8-95.8% viable spores. All quantitative data are shown as mean values plus/minus standard deviation of the mean (SD). Graphs were plotted using GraphPad Prism 10.1.0 (GraphPad software). Statistical analyses were performed using one-way ANOVA with Bonferroni’s post-test correction, or an unpaired t-test. All statistical analyses were performed using Sigmastat 3.5 (SigmaPlot software). *P* values ≥ 0.05 were considered non-significant. NT (no or very few tetrads). ND (not determined).

### Antisera and immunoblotting

The rat anti-HA antibody was purchased from Roche (Basel, Switzerland). The rabbit anti-Hsp104 antiserum was kindly provided by Chung Wang (Academia Sinica, Taiwan). In brief, whole-cell extracts of ∼6 × 10^7^ yeast vegetative cells and sporulating cells precipitated with trichloroacetic acid (TCA, Sigma-Aldrich) were subjected to SDS-PAGE and Western blotting as described previously (48). Hsp104 was used as a loading control. Size in kilodaltons of standard protein markers is labeled to the left of the blots. All vegetative haploid and diploid cells were grown at 30 °C in YPDA medium to an optical density at wavelength 600 nm (OD_600_) of 0.4-0.6. Sporulating cells were grown overnight at 30 °C in YPA medium (1% yeast extract, 2% peptone, and 2% potassium acetate, 0.1% adenine, and 1% tryptophan), then transferred into sporulation medium (3% potassium acetate, 0.02% Raffinose, 0.05% adenine, 1% tryptophan, 0.007% antiform 204) (termed time-point 0), before being harvested at the indicated time-points.

### Determination of the percentages of spore viability, post-MI (%), and sporulation efficiency

Spore viability on the dissection plates was assessed after three days of growth at the indicated temperature (48,82,137). To determine post-MI (%), i.e., the percentage of cells that completed MI and both MI+MII divisions (139,140), sporulation cultures of each cross were harvested 48 hours after being transferred into sporulation medium and then divided into three aliquots. The sporulating cells in the first aliquots (1.0 mL) were collected by centrifugation, resuspended in 1.0 mL of fixation solution (40% ethanol and 0.1M sorbitol) at 4 °C, and then stained with 4, 6-diamidino-2-phenylindole (DAPI). DAPI was excited using a 404 nm laser and emission light was collected with 450/50 BP filters. Fluorescence microscopy images of the stained cells were acquired using a Zeiss Axio Imager Z1 fluorescence microscope and a 63x/1.4 oil immersion objective.

To quantify sporulation efficiency, the second aliquot of sporulation cultures remained untreated, and the third aliquot was stained with DiBAC_4_(5) [Bis-(1,3-dibutylbarbituric acid) pentamethine oxonol], which can stain yeast tetrads and dyads, as well as adhere to spores following bulk tetrad disruption (141). The vital dye DiBAC_4_(5) (Absorption = 590 nm; Emission = 616 nm) is a lipophilic, anionic molecule that is known to fluorescently stain dead yeast cells based on collapse of their membrane potential (142). In brief, sporulating diploid cells were rinsed twice with phosphate-buffered saline (PBS), resuspended in 1 mL PBS, and then stained at room temperature in the dark for 1 minute with 1 μg/mL DiBAC4(5) (Sigma-Aldrich, US) using a 1 mg/mL stock solution in DMSO. After staining, excess dye was removed by rinsing and resuspending the cells in 1 mL PBS. Flow cytometry was performed on a BD FACSymphony A3 Cell Analyzer, as described previously (143). The cells were diluted in PBS to a density of ∼1×10^7^ cells/ml and sorted at room temperature using the instrument’s “ultrapure” setting. Since yeast diploid cells lacking *DMC1* cannot complete MI and hardly form any dyads or tetrads (144), both non-stained samples and DiBAC4(5)-stained *dmc1*Δ sporulating cells were used for gating.

### Genome-wide mapping of interhomolog recombination products generated by wild-type and mutant SK1/S288c hybrid zygotes

We applied Oxford Nanopore Technology (ONT) to sequence genomic DNA. Isolation of genomic DNA, ONT genome sequencing and assembly were carried out as described previously (133,145–147). ONT direct DNA library preparation and sequencing were performed at the Genomics Core of the Institute of Molecular Biology, Academia Sinica. In brief, small fragments of purified genomic DNA were removed by using an Ampure cleanup kit (Beckman Coulter). High-quality DNA quantification was conducted using a Qubit fluorometer (Thermo Fisher Scientific). The average length of genomic DNA was scaled using Femto Pulse (Agilent). We used 1 μg of genomic DNA for library construction. The library contained three or four DNA samples barcoded using the EXP-NBD104 Rapid barcoding kit (Oxford Nanopore Technologies), and adaptors were added with the SQK-LSK109 ligation sequencing kit (Oxford Nanopore Technologies). The library was loaded onto a R9.4.1 Flow Cell system (FLO-MIN106) and processed for 48 hours on the MinION platform. The sequencing process was controlled using MinKNOW software (version 3.6.5; Oxford Nanopore Technologies). All experimental procedures were carried out according to the manufacturer’s instructions. FAST5 data files were generated upon completion of sequencing. These files were converted into FASTQ files in Guppy (v3.6.0). All reads were split into separate FASTQ files based on their barcode by means of the qcat tool (v1.1.0), and then Canu (v2.1.1) (148) was used to perform whole-genome assembly on the corresponding FASTQ files for each sample. The parameter of corOutCoverage was set to 60 based on sequencing read depth. Finally, we used medaka (v1.2.0) to polish the assemblies with the raw reads. The raw datasets have been deposited in the National Center for Biotechnology Information (Bioproject accession number PRJNA1216236). Since Canu generated expected numbers of contigs for each sample, and a comparison of the genome to wild-type S288c indicated no broken contigs, we did not perform any manual finishing or validation. Genome-wide mapping of interhomolog recombination products was performed using the same approaches described in our previous recombination analyses of *Trichoderma reesei* and *S. cerevisiae* hybrid meiosis (5,132,133).

### Physical analysis by Southern hybridization of meiotic recombination

Synchronous meiosis was performed as described previously (139,140). Genomic DNA extraction and 1D gel analysis were also performed as described previously (55,136). The Southern blotting physical analysis was carried out using “Probe A”, as described previously (36). DNA probes were radiolabeled using a Random Priming kit (Invitrogen, USA) (56). Hybridizing species were quantitated using a Fuji BAS2000 phosphorimager and Image Gauge V3.3 software (Fuji Photo Film Co. Ltd.) (149).

### Analysis of CO frequencies and interferences

Diploids were sporulated in liquid sporulation medium and the recombination between fluorescence markers on chromosome VIII was scored after 48 hours of sporulation using a Zeiss Axio Imager Z1 fluorescence microscope, as described previously (138). Two independent sets of each strain were combined and 977-3504 tetrads were scored for COs in two test intervals and for MI NDJ events. Genetic distances in the *CEN8-ARG4* and *ARG4-THR1* intervals were calculated from the distribution of parental ditype (PD), nonparental ditype (NPD), and tetratype (T) tetrads, and genetic distances (cM) were calculated using Perkins equation: cM (centimorgan) = [100 (6NPD + T)] / [2 (PD + NPD + T)]. Standard errors of genetic distances were calculated using Stahl Laboratory Online Tools (https://elizabethhousworth.com/StahlLabOnlineTools) (150). Interference (γ) is defined as γ = 1 – (coefficient of coincidence or c.o.c.), where c.o.c. is calculated by dividing the actual frequency of double recombinants by the expected frequency (74). Both genetic distances and the “NPD ratio” (*fN_obs_*/*fN_exp_*) were also calculated using Stahl Laboratory Online Tools. The NPD ratio is an indication of the intensity of interference, with smaller values indicating stronger positive interference (151,152).

### Statistical analysis

We combined three independent sets of each strain and scored tetrads either individually (N/3; split in triplicate) or collectively (i.e., N; total). All quantitative data is shown as mean plus/minus Standard Deviation of the mean (SD), as indicated in each figure legend. Graphs were plotted using GraphPad Prism 10.1.0 (GraphPad software). Statistical analyses were performed using two-way analysis of variance (ANOVA, for comparing genetic distances of wild-type purebred and hybrid strains), one-way ANOVA (for comparing spore viability, γ, and MI NDJ of wild-type purebred and hybrid strains) with Bonferroni’s post-test correction or an unpaired t-test. All statistical analyses were performed using Sigmastat 3.5 (SigmaPlot software). *P* values ≥ 0.05 were considered non-significant.

## Results

### Identification of anti-MMR genes preferentially required for intraspecies hybrid meiosis rather than for purebred meiosis

We established a screen for *S. cerevisiae* anti-MMR genes using three different wild-type diploid strains (SK1/SK1-*HL* purebred zygotes, and SK1/S288c-*HL*\* and SK1/S288c-*HL*\**-SNP+* hybrid zygotes; see details below) (Figure 1A and DS18). In brief, all diploid cells were incubated at 30 °C in sporulation medium for at least 48 hours and then scored for their capability to form mature tetrads and viable sexual spores. Spore viability was determined by isolating the four single spores from each tetrad on YPDA plates. The SK1/SK1-*HL* purebred strains carry a well-characterized DSB and meiotic recombination hotspot, i.e., *HIS4::LEU2-*(*NBamHI*)*/his4-X::LEU2-*(*NgoMIV*)*- URA3* (referred to herein as “*HL*”) (Figure 1A and DS18) (51,56,153). *HL* constitutes a recombination reporter in which DSBs form at high frequency within a narrow zone of ∼150 base pairs (bp), making this hotspot suited for genetic analysis, as well as for physical analyses of DSB resection and the subsequent choice of DSB repair via the IH-HR or inter-sister (IS-HR) pathways (35,56,91). We applied polymerase chain reaction (PCR) to amplify a DNA fragment containing the *HIS4::LEU2-*(*New BamHI*) allele from the SK1-*HL*(*MATα*) haploid strain and then integrated it into a haploid S288c(*MATα*) strain. The resulting SK1/S288c-*HL*\* hybrid diploid cells carry the same *HL* DSB hotspot, except that a point mutation (*) was introduced to mutate the *XhoI* restriction site adjacent to probe A (Figure 1A). To investigate the impacts of denser mismatched base pairs in hDNA during DSB repair, we also generated a SK1/S288c-*HL*\**-SNP+* hybrid strain that carries the *HIS4::LEU2-*(*New BamHI*)*-SNP+/his4-X::LEU2-*(*NgoMIV*)*-URA3* hotspot. Compared to the *HIS4::LEU2-*(*New BamHI)* allele, the *HIS4::LEU2-*(*New BamHI*)*-SNP+* allele possesses 86 exogenously introduced single nucleotide polymorphisms (SNPs; indicated in green, Figure 1A), as it was modified to harbor single nucleotide sequence change every 30.5 ± 12.2 bp for 1,633 bp to the left of the DSB site and 944 bp to the right. The nucleotide sequences of four loci (∼13,000 bps) shown in Figure 1A are provided in DS19. Southern hybridization revealed that all three of these diploid strains are feasible for quantification of IH-CO and IH-NCO recombination products (see below). A similar approach was used previously to study repeated strand invasion and extensive branch migration in *S. cerevisiae* meiosis (50).

First, we confirmed that *rad54*Δ (154) and *dmc*1Δ *hed*1Δ (45) resulted in fewer viable spores in the two hybrid zygotes (i.e., SK1/S288c-*HL*\* and SK1/S288c-*HL*\**-SNP+*) compared to the purebred zygote (i.e., SK1/SK1-*HL*). Deletion of *RAD51* (*rad51*Δ) resulted in mature tetrads but no viable spores in the SK1/SK1-*HL* purebred zygotes, as well as in the SK1/S288c-*HL\** and SK1/S288c-*HL*-SNP+* hybrid zygotes. Moreover, *rad51-II3A* (155) and *rad51-*Δ*N* (48) also led to lower fertility in the SK1/S288c-*HL\** and SK1/S288c-*HL*-SNP+* hybrid zygotes relative to the SK1/SK1-*HL* purebred zygotes (Figure 1B and DS2). As reported previously (155), *rad51-II3A* hardly affected the meiotic recombination or spore viability of the SK1/SK1-*HL* purebred zygotes. The Rad51-II3A mutant protein has a normal DNA binding site I but a defective DNA binding site II. The two binding sites in Rad51 and Dmc1 are referred to herein as Rad51-I, Rad51-II, Dmc1-I, and Dmc1-II, respectively. Therefore, Rad51-II3A mutant protein can assemble onto ssDNA and form nucleoprotein filaments, but it is defective in capturing a homologous dsDNA template for homology pairing and strand exchange (155). We also observed that *rad54*Δ *dmc*1Δ *hed*1Δ SK1/SK1 purebred zygotes hardly produced mature tetrads, indicating that Rad54 is indispensable for Rad51-mediated IH-HR in the *dmc*1Δ *hed*1Δ line (Figure 1B and DS20).

Next, we screened ∼50 other mutants (Table S1) of proteins known to be involved in DSB repair and/or MMR in *S. cerevisiae* meiosis. Their genotypes and fertility phenotypes (i.e., ability to form tetrads and/or viable spores) are provided in DS18 and DS20, respectively. We classified these mutants into five distinct groups based on their ability to form tetrads and viable spores in the SK1/SK1-*HL* purebred zygotes, as well as in both wild-type and *P_CLB2_-MSH2* SK1/S288c-*HL*\**-SNP+* hybrid zygotes, respectively (Figure 2).

**Figure 2.**
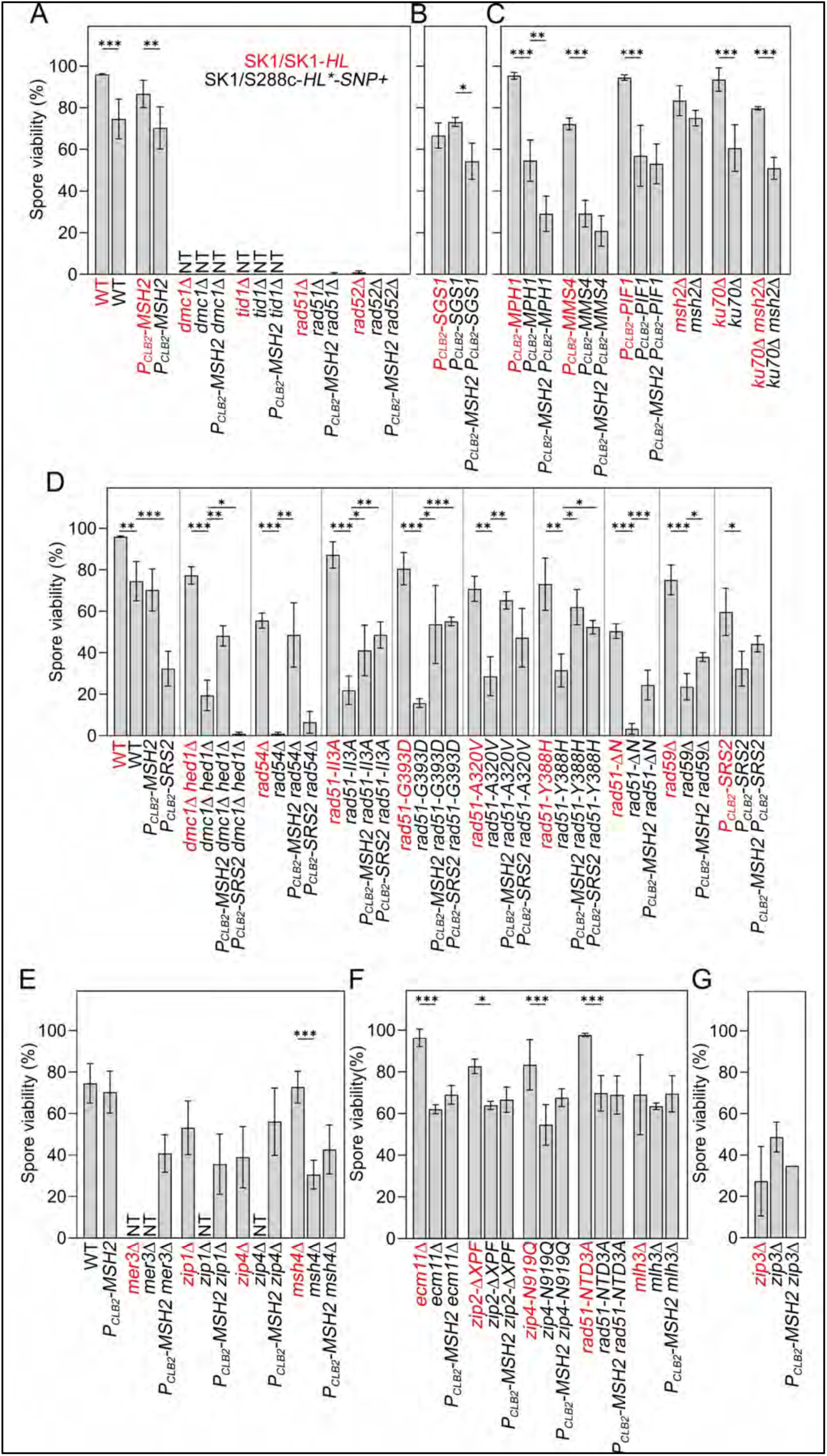
Classification of yeast genes with or without anti-MMR activities. Spore viability of six different types of yeast mutants: type I (**A**), type II (**B**), type III (**C**), type IVA (**D, E**), type IVB (**F**), and type V (**G**). The anti-MMR (type IVA) mutants (except *mer3*Δ) resulted in lower fertility (spore viability) in hybrid zygotes compared to purebred zygotes. The low fertility phenotype of S288c/SK1 hybrid zygotes with anti-MMR mutant(s) could be rescued by *P_CLB2_-MSH2.* NT (no or very few tetrads). All statistical analyses were carried out as described in Figure 1.

The group I mutants include *rad51*Δ, *rad52*Δ*, dmc1*Δ, and *tid1*Δ. Both purebred and hybrid zygotes hosting these mutants formed very few tetrads and/or viable spores and were insensitive to the introduction of *P_CLB2_-MSH2* (Figure 2A and DS20).

The group II mutants (*P_CLB2_-SGS1*, *sgs1*Δ*-C795* and *sgs1*Δ) produced similar numbers of viable spores in the SK1/SK1-*HL* purebred zygotes (66.7±6.1%, 73.6±6.1%, and 44.0±4.2%) and in the SK1/S288c-*HL*-SNP+* hybrid zygotes (73.1±2.1%, 66.2±4.2%, and 53.2±4.2%), respectively. Introduction of *P_CLB2_-MSH2* into the *P_CLB2_-SGS1* SK1/S288c-*HL*-SNP+* hybrid zygotes slightly reduced the spore viability from 73.1±2.1% to 54.4±8.6% (Figure 2B and DS20). Thus, Sgs1 does not meet the criteria for an anti-MMR protein in our assay.

The group III mutants (*P_CLB2_-MPH1*, *P_CLB2_-MMS4* and *P_CLB2_-PIF1*) produced more viable spores in the SK1/SK1-*HL* purebred zygotes than in the SK1/S288c-*HL*-SNP+* hybrid zygotes and *P_CLB2_-MSH2* SK1/S288c-*HL*-SNP+* hybrid zygotes. *ku70*Δ also belongs to this group. The order of their spore viability was: *ku70*Δ SK1/SK1-*HL* > *ku70*Δ *msh2*Δ SK1/SK1-*HL* > *ku70*Δ SK1/S288c-*HL*-SNP+* > *ku70*Δ *msh2*Δ SK1/S288c-*HL*-SNP+* (Figure 2C and DS20). We conclude that Mph1, Mms4, Pif1, and the Ku70/80 complex do not meet the criteria for anti-MMR proteins.

The group IV mutants are “anti-MMR” mutants, which we divided into two subgroups, IV-A (Figure 2D-2E) and IV-B (Figure 2F), according to the degree of hybrid infertility. Except for *mer3*Δ, all other “IV-A” mutants showed more pronounced fertility (spore viability) defects in the SK1/S288c-*HL*-SNP+* hybrid zygotes than in the SK1/SK1-*HL* purebred zygotes, including *dmc1*Δ *hed1*Δ, *rad54*Δ, *rad51-II3A*, *rad51*-Δ*N*, *rad51-A320V*, *rad51-Y388H*, *rad51-G393D*, *rad59*Δ, *P_CLB2_-SRS2*, *mer3*Δ, *zip1*Δ, *zip4*Δ, and *msh4*Δ. The low spore viability defects of all group IV-A mutants (including *mer3*Δ) in the SK1/S288c-*HL*-SNP+* hybrid zygotes could be rescued (at least in part) by introducing *P_CLB2_-MSH2* (Figures 2D-2E). We also found that *P_CLB2_-SRS2* can increase fertility of the SK1/S288c-*HL*-SNP+* hybrid zygotes with *rad51-II3A*, *rad51*-Δ*N*, *rad51-A320V*, *rad51-Y388H*, or *rad51-G393D*, but not those with *dmc1*Δ *hed1*Δ or *rad54*Δ (Figure 2D). The “IV-B” mutants exhibited much weaker or no anti-MMR activities, including *ecm11*Δ, *zip2-ΔXPF*, *zip4-N919Q*, *rad51-NTD3A*, and *mlh3*Δ (Figure 2F).

The only group V mutant is *zip3*Δ, which resulted in more viable spores in the *MSH2* SK1/S288c-*HL*-SNP+* hybrid zygotes (48.6±7.2%) than in *MSH2* SK1/SK1-*HL* purebred zygotes (27.3±16.7%) and *P_CLB2_-MSH2* SK1/S288c-*HL*-SNP+* hybrid zygotes (34.7±0%) (Figure 2G).

We then confirmed by immunoblotting experiments using anti-HA antibodies that *P_CLB2_* can diminish the steady-state protein levels of HA-tagged Msh2, Sgs1, and Srs2 in sporulating cells, but not in vegetative cells, respectively (DS21 and Supplementary Figure S1).

These genetic results reveal several intriguing and important features of the anti-MMR proteins. First, the anti-MMR function of Rad51 requires a sufficient amount of catalytically active Rad51 (as revealed by Rad51-ΔN experiments) (48), Rad51’s binding capability to Rad52 and/or Srs2 (as revealed by Rad51-A320V, Rad51-Y388H, and Rad51-G393D) (27), the homology pairing and strand exchange activities of Rad51 itself (as revealed by Rad51-II3A) (155), and Rad51’s homology pairing and strand exchange cofactor Rad54 (154,156,157). In contrast, the anti-MMR activity of Rad51 is hardly affected by Mec1/Tel1-dependent NTD phosphorylation. Wild-type Rad51, Rad51-NTD3A, and Rad51-ΔN are all catalytically active in homology-directed DSB repair during meiosis, with *rad51*Δ, *rad51-*Δ*N*, and *rad51-NTD3A* meiotic cells producing <1%, ∼50% and ∼97% viable spores, and the steady-levels of Rad51 proteins in meiotic cells being in the order of: wild-type ∼ Rad51-NTD3A >> Rad51-ΔN (48). Second, the group of ZMM proteins responds differentially to genetic polymorphism and MMR in hybrid zygotes. Mer3, like Dmc1 and Tid1, is indispensable for purebred and hybrid zygotes to sporulate and form viable spores. The anti-MMR activities of Mer3, Zip1, and the ZZS complex are significantly stronger than those of the MutSγ complex, the MutLγ complex, or the Ecm11-Gmc2 complex. Moreover, the anti-MMR activities of the ZZS complex are not significantly affected by loss of the Zip4-Ecm11 interaction (89) or the Zip2-Spo16 interaction (90,91). In contrast, Zip3 may function independently of or in weak coordination with the MMR system in hybrid zygotes.

### Srs2, like *E. coli* UvrD, facilitates the MMR system in antagonizing formation of Rad51-mediated D-loops

Both *P_CLB2_-MSH2* and *P_CLB2_-SRS2* increased the fertility of the SK1/S288c-*HL*-SNP+* hybrid zygotes harboring *rad51-II3A*, *rad51*-Δ*N*, *rad51-A320V*, *rad51-Y388H*, or *rad51-G393D.* In contrast, only *P_CLB2_-MSH2*, but not *P_CLB2_-SRS2*, increased the fertility of the SK1/S288c-*HL*- SNP+* hybrid zygotes with *dmc1*Δ *hed1*Δ or *rad54*Δ (Figure 2D). Srs2 is the *S. cerevisiae* homolog of *E. coli* MMR helicase UvrD. UvrD can dismantle RecA nucleoprotein filaments during the process of nucleotide excision repair and methyl-directed MMR (72), whereas the main function of Srs2 in HR is to dissemble the Rad51-ssDNA nucleoprotein filaments (22). Accordingly, Srs2 and UvrD not only are highly conserved in their amino acid sequences and protein structures, but also exhibit anti-Rad51 and anti-RecA activities to facilitate the MMR system in *S. cerevisiae* and *E. coli*, respectively.

### The anti-MMR activities of Rad51 and Rad54 acts during and after the formation of Rad51-made D-loops

We propose that the Rad51/Rad54 recombinasome and the MMR/Srs2 ensemble are yin-yang partners that antagonize each other during Phase II of meiosis. The anti-MMR activities of Rad51 and Rad54 outweigh their pro-HR functions when more mismatched base pairs in hDNA are generated at or near DSBs with sequence heterozygosity in hybrid zygotes, as well as at the *HIS4::LEU2/his4-X::LEU2-URA3* hotspot in purebred zygotes (see below). The anti-MMR activity of Rad51 acts during Rad51-mediated D-loop formation to antagonize Srs2 and MMR in hybrid zygotes, whereas Rad54 acts later to prevent the Msh2-containing complexes from binding to the mismatched base pairs in the emerging D-loop made by Rad51 or those in the resulting joint molecules. This hypothesis is consistent with our results presented herein and those of several previous reports. First, Dmc1 is a potent inhibitor of Srs2 (20). Second, Rad54 binds tightly to Rad51-ssDNA and acts downstream of Srs2 (22). Third, Rad54 not only possesses chromatin remodeling activity to clear nucleosomes or other proteins during homology pairing (23), but also exhibits Rad51-dependent branch migration activity to dissociate Rad51-generated D-loops but not Dmc1-made D-loops (24,25). The anti-MMR activity of Rad54 likely operates through its chromatin remodeling and branch migration functions, separating the Msh2-containing complexes from mismatched base pairs in Rad51-made D-loops and/or joint molecules. Fourth, Rad54 is essential for Rad51-mediated IH-HR in *dmc1*Δ *hed1*Δ purebred zygotes (Figure 1B). However, due to chronic overactivation of the DNA damage checkpoint in the *dmc1*Δ *hed1*Δ meiotic cells (48), Mek1-dependent Rad54 phosphorylation negatively regulates Rad51 activity (46). Since *dmc1*Δ *hed1*Δ purebred zygotes produce more viable spores than *dmc1*Δ *hed1*Δ hybrid zygotes, the anti-MMR function of Rad54 is more important than its pro-HR function in the *dmc1*Δ *hed1*Δ hybrid zygotes. Accordingly, only *P_CLB2_-MSH2* but not *P_CLB2_-SRS2* could rescue the infertility phenotypes of *rad54*Δ and *dmc1*Δ *hed1*Δ hybrid zygotes (Figure 2D).

The results of our DiBAC_4_(5) (Figure S2A) and DAPI staining experiments (Figure S2B-S2D) also revealed that both purebred and hybrid zygotes with *rad51-II3A, rad51-G393D, rad54*Δ, *P_CLB2_-MSH2, rad51-II3A P_CLB2_-MSH2, rad51-G393D P_CLB2_-MSH2, rad54*Δ *P_CLB2_-MSH2*, *P_CLB2_-SRS2, P_CLB2_-SRS2 rad51-II3A*, or *P_CLB2_-SRS2 rad54*Δ can sporulate and complete MI. In contrast, the *dmc1*Δ *hed1*Δ hybrid zygotes exhibit lower sporulation efficiency than *dmc1*Δ *hed1*Δ purebred zygotes and *dmc1*Δ *hed1*Δ *P_CLB2_-MSH2* hybrid zygotes (Figure S2A). Moreover, *dmc1*Δ *hed1*Δ also delayed MI progression (Figures S2C-S2D).

### The anti-MMR activity of Rad51 preferentially targets MutSα, MutSα, and MutLα rather than MutLα or MutLγ

Next, we observed that, in the SK1/S288c-*HL*-SNP+* hybrid zygotes, the order of spore viability was *P_CLB2_-MSH6* ∼ *P_CLB2_-MSH3* ∼ *msh2*Δ ∼ *mlh2*Δ ∼ *P_CLB2_-MSH6 rad51-II3A* ∼ *P_CLB2_-MSH2 ∼ mlh3*Δ *∼ pms1*Δ ∼ *pms1*Δ *rad51-II3A* ∼ *P_CLB2_-MSH3 rad51-II3A* ∼ *msh2*Δ *rad51-II3A* ∼ *P_CLB2_- MSH2 rad51-II3A* > *mlh2*Δ *rad51-II3A* > *rad51-II3A* > *mlh3*Δ *rad51-II3A* (Figure 3 and DS20). We conclude that the anti-MMR activity of Rad51 can act on MutS-driven mismatch recognition and MutLα-mediated mismatch correction, but not MutLγ-dependent IH-CO. Our results also indicate that *P_CLB2_-MSH6* was slightly better than *msh2Δ, P_CLB2_-MSH2,* or *P_CLB2_-MSH3* in rescuing the infertility of *rad51-II3A* hybrid zygotes. One possibility is that mutations in *MSH2* cause a strong general mutator phenotype, resulting in an accumulation of both frameshift and single-base substitute mutations, high rates of frameshift mutation reversion, and dinucleotide repeat instability. In contrast, mutations in *MSH6* cause a modest mutator phenotype that is confined primarily to an accumulation of single-base substitute mutations (158).

**Figure 3.**
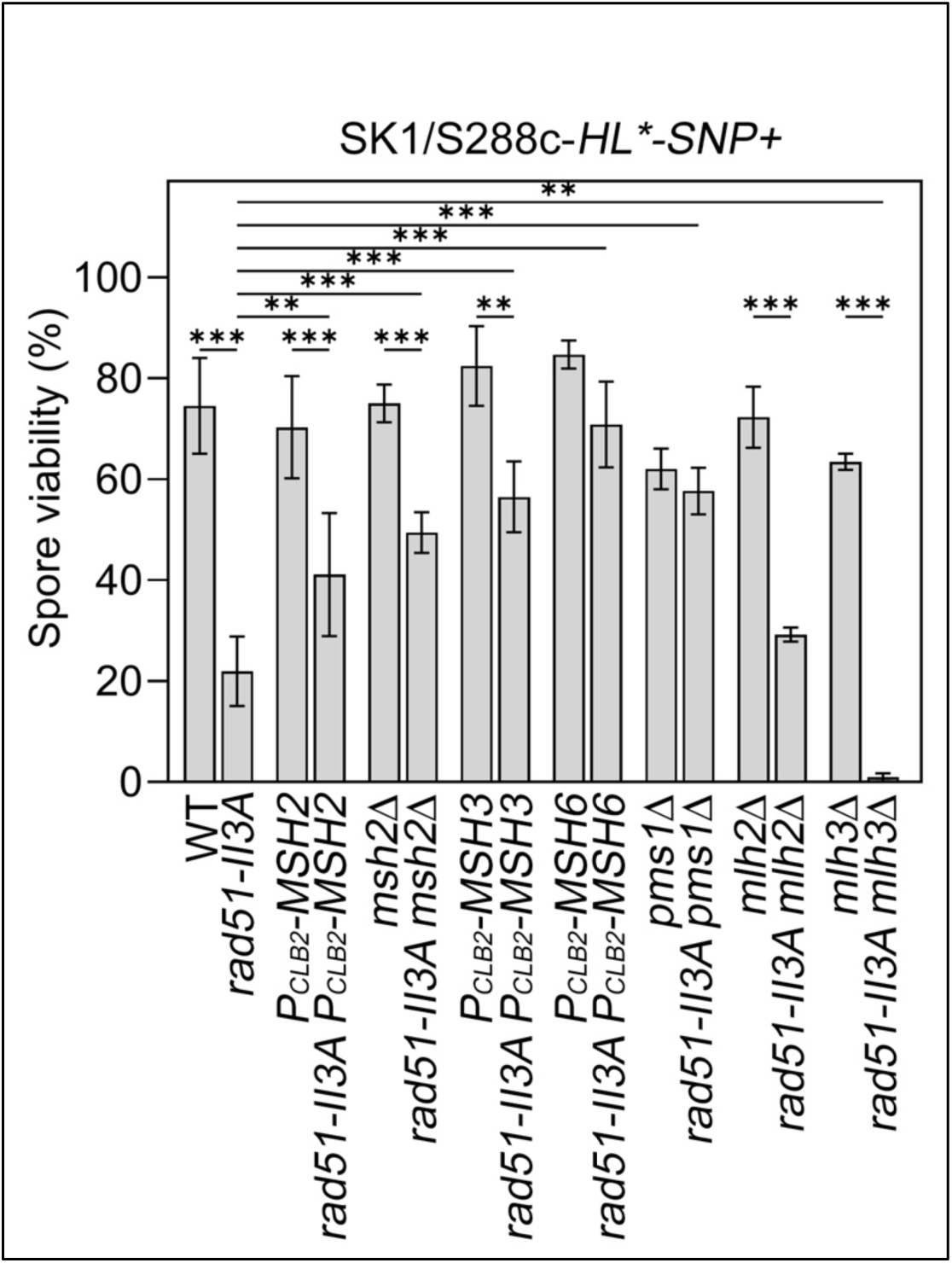
Rad51 preferentially antagonizes MutSα, MutSα, and MutLα rather than MutLα or MutLγ. Spore viability of *rad51-II3A* hybrid zygotes with indicated MMR mutations. All statistical analyses were carried out as described in Figure 1.

### Rad51 and Dmc1 utilize different mechanisms to encounter mismatched base pairs and the MMR system during homology-directed DSB repair

Controversy exists regarding the roles of Rad51 and Dmc1 in tolerating mismatched base pairs during the homology pairing and strand exchange reactions. First, *S. cerevisiae* Rad51 and Dmc1 exhibit similar mismatch tolerability during homology pairing strand exchange *in vivo* (159). Second, Dmc1 is superior to Rad51 in tolerating the mismatched base pairs in hDNA in budding yeast, fission yeast, and mammals (5,45,160–165). Two hypothetical cryogenic electron microscopy (cryo-EM) protein models have been used to propose that two critical amino acids in eukaryotic Rad51 (i.e., *H. sapiens* Val273 and Asp274, *S. cerevisiae* Val331 and Asp332) and Dmc1 (*H. sapiens* Pro274 and Gly275, *S. cerevisiae* Pro267 and Gly268) might explain why Dmc1 exhibits higher mismatch tolerability than Rad51 (164,165) (Table S1 and DS20). Our genetic results do not support this hypothesis because the capability of wild-type Rad51 in generating viable spores in both SK1/SK1 *dmc1*Δ *hed1*Δ purebred zygotes and SK1/S288c hybrid *dmc1*Δ *hed1*Δ zygotes is similar to that of Rad51-V331P, Rad51-D332G or the Rad51-V331P D332G double mutant. Moreover, Pro267 is dispensable for Dmc1’s function in meiosis because both *dmc1-P267V* purebred and hybrid zygotes (96.8% and 84.3%) produced as many viable spores as purebred and hybrid wild-type zygotes (96.4% and 85.1%). In contrast, Gly268 is essential for Dmc1’s catalytic function. Like *dmc1*Δ, both *dmc1-G268D* and the *dmc1-P267V dmc1-G268D* double mutant elicited severe meiotic arrest before MI and resulted in no tetrads being generated in SK1/SK1 purebred and SK1/S288c hybrid zygotes (DS20).

We propose that Rad51 and Dmc1 utilize different mechanisms to tolerate mismatched base pairs in hDNA and to antagonize the MMR system during *S. cerevisiae* hybrid meiosis. In Phase I, Rad51 is inhibited by Hed1 (32). Since Dmc1 and Tid1 (Rdh54) do not possess anti-MMR activities (Figure 2A) and Dmc1 exhibits higher mismatch tolerability than Rad51 (5,45,160–165), the Dmc1-mediated IH-HR pathway likely adopts two different approaches to tolerate mismatched base pairs and antagonize MMR, respectively. Before mismatched base pairs are recognized by the Msh2-containing MMR complexes, they are likely tolerated by Dmc1-ssDNA nucleoprotein filaments, thereby initiating Dmc1-mediated D-loop formation and also promoting SC assembly or homology synapsis (32). This early function of Dmc1 requires Tid1 (21), Rad51-I (i.e., Rad51’s capability to bind ssDNA), Rad52, Dmc1-I, Dmc1-II (i.e., Dmc1’s capability to capture dsDNA during the homology pairing and strand exchange reactions), but not Rad51-II (155). Accordingly, *dmc1*Δ*, tid1*Δ, *rad51*Δ, and *rad52*Δ all result in strong infertility defects in both SK1/SK1 purebred zygotes and SK1/S288c hybrid zygotes. Introduction of *P_CLB2_-MSH2* into the SK1/S288c-*HL*- SNP+* hybrid zygotes also cannot rescue the infertility defects of the *dmc1*Δ*, tid1*Δ, *rad51*Δ, and *rad52*Δ mutants (Figure 2A and DS20).

### ZMM proteins antagonize MMR and Sgs1 to facilitate Dmc1- and Rad51-mediated IH-CO formation

Dmc1 repairs most DSBs in Phase I of meiosis, whereas Rad51 repairs residual DSBs in Phase II (32). After the formation of Dmc1-generated D-loops in WT Phase I and the formation of Rad51-made D-loops in Phase II of WT and *dmc1*Δ *hed1*Δ cells, the Msh2-containing MMR complexes recognize the mismatched base pairs initially present in Dmc1- and Rad51-generated D-loops, as well as the mismatched base pairs in the resulting joint molecules generated during DNA repair synthesis, branch migration, or multiple rounds of strand invasion. Sgs1 acts downstream of Msh2-mediated mismatch recognition (127) and it promotes IH-NCO formation (54). Sgs1 may also function in an MMR-independent manner during IS-HR and IH-HR in homozygous meiosis (i.e. between identical maternal and paternal genomic sequences). Both scenarios explain why IH-NCO is formed earlier than IH-CO during *S. cerevisiae* meiosis (51–53). Subsequently, Dmc1-made joint molecules in Phase I are better than Rad51-made joint molecules in Phase II in recruiting Mer3 (52,65) and other ZMM proteins (Zip1, the ZZS complex, and MutSγ). These synapsis-promoting ZMM proteins not only induce class I IH-CO formation, but also antagonize MMR and Sgs1 (Figure 2E), thereby preventing excessive IS-NCO and IH-NCO formation. Since *rad51-II3A* hybrid zygotes with *mer3*Δ, *zip1*Δ, *zip3*Δ, or *zip4*Δ do not form mature tetrads and *rad51-II3A* hybrid zygotes with *mlh3*Δ or *msh4*Δ can form a few mature tetrads but very few viable spores (Figure 4A), Dmc1- and Rad51-mediated IH-CO pathways are both significantly diminished in *rad51-II3A zmm*Δ hybrid zygotes (see below).

**Figure 4.**
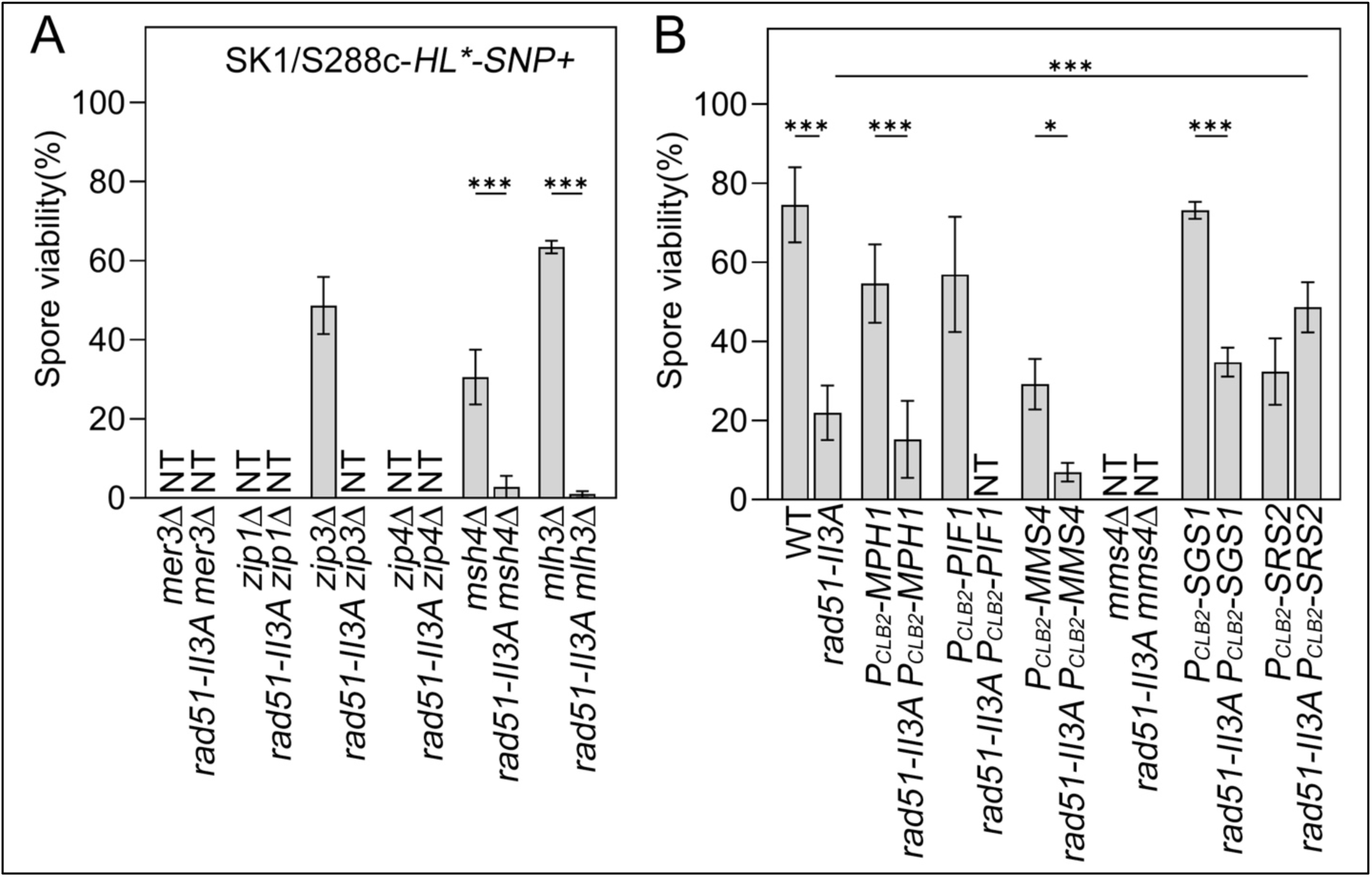
The effects of *zmm*Δ (A), *P_CLB2_-MPH1, P_CLB2_-PIF1, P_CLB2_-MMS4, mms4*Δ, *P_CLB2_-SGS1* or *P_CLB2_-SRS2* (B) on fertility of WT and *rad51-II3A* hybrid zygotes. NT (no or very few tetrads). All statistical analyses were carried out as described in Figure 1.

Srs2 and Sgs1 respond differently to mismatched base pairs and the MMR system. Suppression of recombination between homeologous sequences was found to be enhanced in *srs2* mutants but reduced in *sgs1* mutants (166). We confirmed that *P_CLB2_-MSH2* has the opposite effect to *P_CLB2_-SRS2* and *P_CLB2_-SGS1* in the SK1/S288c-*HL*-SNP+* hybrid zygotes, with the order of spore viability being *P_CLB2_-SRS2* (32.4%) < *P_CLB2_-MSH2 P_CLB2_-SRS2* (44.3%) < *P_CLB2_-MSH2 P_CLB2_-SGS1* (54.4%) < *P_CLB2_-MSH2* (70.3%) ∼ *P_CLB2_-SGS1* (73.1%) (Figures 2A, 2B, 2D, and DS2). *P_CLB2_-SRS2* proved slightly better than *P_CLB2_-SGS1* in rescuing the low fertility phenotype of *rad51-II3A* SK1/S288c-*HL*-SNP+* hybrid zygotes, with the order of spore viability being *rad51-II3A* (21.9%) < *rad51-II3A P_CLB2_-SGS1* (34.7%) ∼ *rad51-II3A P_CLB2_-MSH2* (41.1%) < *rad51-II3A P_CLB2_-SRS2* (48.6%) (Figure 4B).

Notably, *P_CLB2_-MSH2, P_CLB2_-SGS1*, *P_CLB2_-MSH2 P_CLB2_-SGS1,* or *P_CLB2_-MSH2 P_CLB2_-SRS2* increased the fertility of the *zip4*Δ SK1/S288c-*HL*-SNP+* hybrid zygotes from generating no mature tetrads to 56.0%, 54.2%, 67.6%, and 31.0% viable spores, respectively. In contrast, *P_CLB2_-SRS2* did not rescue the infertility of the *zip4*Δ SK1/S288c-*HL*-SNP+* hybrid zygotes (Figure 5).

**Figure 5.**
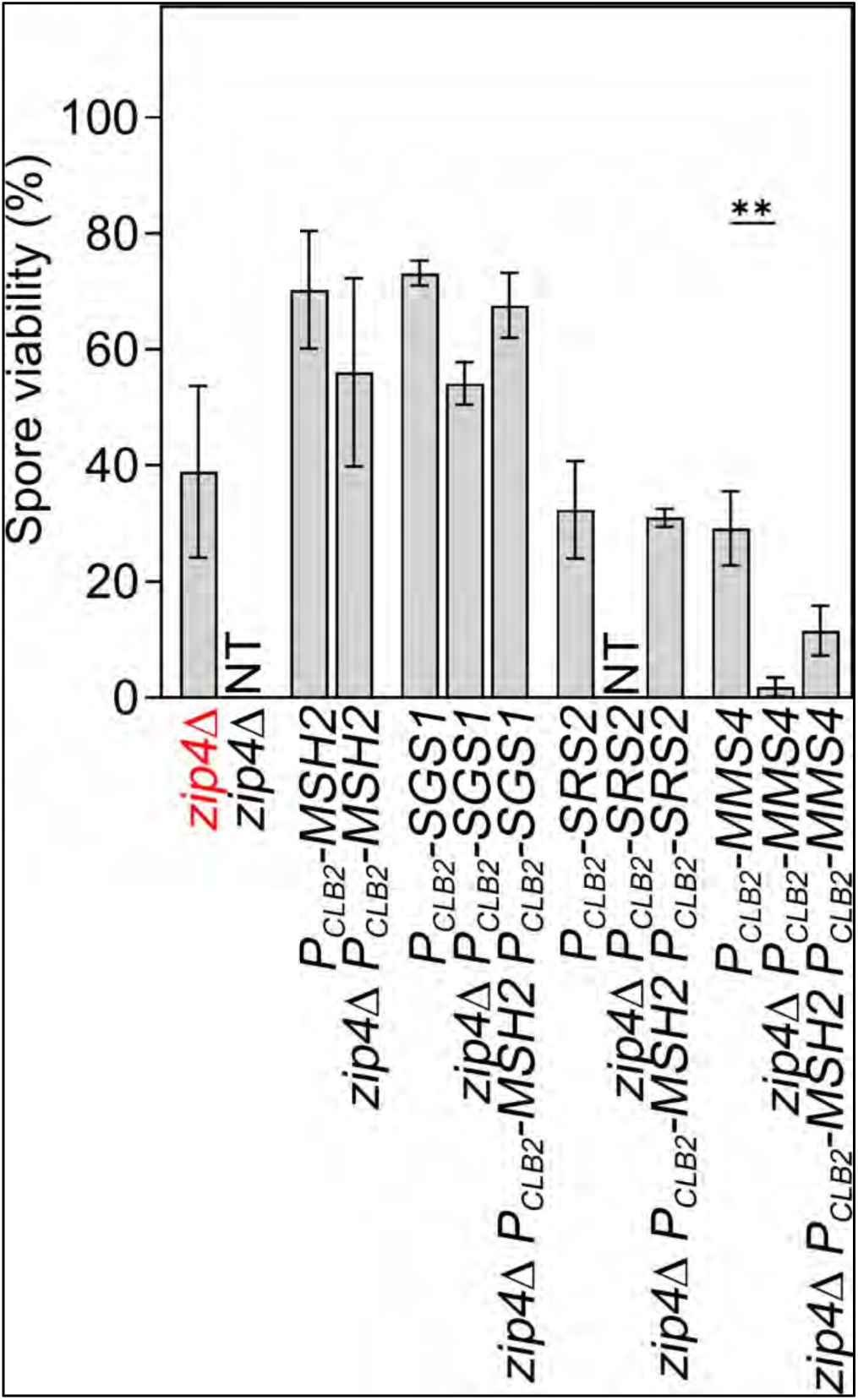
The effects of *P_CLB2_-SGS1, P_CLB2_-SRS2,* or *P_CLB2_-MMS4* on fertility of WT, *zip4*Δ and *zip4*Δ *P_CLB2_-MSH2* hybrid zygotes. SK1/SK1-*HL* purebred zygotes (red); SK1/S288c-*HL*- SNP+* hybrid zygotes (black). NT (no or very few tetrads). All statistical analyses were carried out as described in Figure 1.

In conclusion, Srs2 disassembles the Rad51-ssDNA filaments, thus antagonizing Rad51’s anti-MMR function during Rad51-mediated D-loop formation in hybrid zygotes. After the formation of Rad51-made D-loops, Rad54’s chromatin remodeling activity (23) and/or branch migration activity (24,25) prevents (or even removes) the Msh2-containting MMR complexes from associating with the mismatched base pairs in the Rad51-generated D-loops and the resulting joint molecules. The pro-HR functions of Mer3, Zip1, the ZZS complex, and the MutSγ complex promote both Dmc1-made joint molecules and Rad51-made joint molecules to form class I IH-COs. The anti-MMR activities of ZMM proteins also prevent the Msh2-containing MMR complexes from associating with the mismatched base pairs in both Dmc1- and Rad51-mediated joint molecules. Sgs1 acts downstream of Msh2-mediated mismatch recognition during homeologous recombination (127). Accordingly, the anti-MMR activities of ZMM proteins prevent Sgs1-mediated IH-NCO formation in both Phase I and Phase II, whereas Rad54’s anti-MMR activity only prevents Sgs1-mediated IH-NCO formation in Phase II.

### Possible roles of Mph1, Pif1 and Mms4 in hybrid meiosis

Mph1, Pif1, and Mms4•Mus81 do not exhibit anti-MMR activities, but they are functionally more important in hybrid zygotes than in purebred zygotes (Figure 2C). *P_CLB2_-MPH1*, *P_CLB2_-PIF1, P_CLB2_-MMS4,* or *mms4*Δ further reduced the fertility of *rad51-II3A* SK1/S288c-*HL*-SNP+* hybrid zygotes (Figure 4B). Therefore, Mph1, Pif1, and Mms4•Mus81, like ZMM proteins, play important roles in promoting both Dmc1- and Rad51-mediated IH-HR during hybrid meiosis.

Mph1 can dissociate Rad51-made D-loops during vegetative growth (30). During meiosis, Mph1 ensures high levels of IH-HR products between non-sister homologous chromosomes in purebred zygotes by preventing precocious DSB strand exchange between sister chromatids before homologs have completed pairing (31). Zip1 antagonizes Mph1-mediated dissociation of late-occurring joint molecules (31). Since *P_CLB2_-MPH1* purebred zygotes and hybrid zygotes generate 95.4% and 54.6% viable spores (Figure 2C), whereas *P_CLB2_-MPH1 P_CLB2_-MSH2* hybrid zygotes only produce 29.2% viable spores, we suggest that the DNA helicase activity of Mph1 may be positively regulated by binding of Msh2-containing MMR complexes to mismatched base pairs in hybrid zygotes. This hypothesis is consistent with a previous report that Mph1 is intricately associated with MutSα during homeologous recombination repair and that MutSα specifically suppresses the formation of COs by inhibiting double Holliday Junction (HJ) formation (167). Separation-of-function mutants of Mph1 defective in Mph1-MutSα association are needed to test this hypothesis.

Although Pif1 was shown previously to specifically promote Rad51-mediated IH-HR but not Dmc1-mediated IH-HR in purebred zygotes (32), two lines of evidence indicate that Pif1 may also have important roles in promoting IH-CO in hybrid zygotes. First, *P_CLB2_-PIF1* purebred zygotes and hybrid zygotes generate 94.4% and 56.9% viable spores (Figure 2C). Second, *rad51-II3A* purebred and hybrid zygotes generate 87.2% and 21.9% viable spores, respectively. *P_CLB2_-PIF1 rad51-II3A* purebred zygotes generate 69.4% viable spores. In contrast, *P_CLB2_-PIF1 rad51-II3A* hybrid zygotes hardly form mature tetrads (Figure 4B and DS20). Pif1 may complete its novel task in hybrid zygotes by limiting the IH-NCO formation mediated by Sgs1. First, Pif1 counteracts Sgs1 and that this genetic interaction between Pif1 and Sgs1 is dependent upon HR and DNA topoisomerase III (Top3) (168). Second, Pif1’s activity that promotes meiotic DNA repair synthesis is restrained by the Mer3-MutLβ ensemble which in turn prevents long gene conversion tracts and possibly associated mutagenesis (93).

The class II IH-CO pathway involves Mms4•Mus81 (85) and, to a lesser degree, Yen1 and Slx1•Slx4 (100,101). Mms4•Mus81 does not possess anti-MMR activity (Figure 2C). Although *mms4*Δ and *mms4*Δ *rad51-II3A* hybrid zygotes cannot form mature tetrads, 50-60% *mms4*Δ and *mms4*Δ *rad51-II3A* hybrid zygotes can still sporulate and complete MI. In contrast, wild-type and *rad51-II3A* hybrid zygotes with *mer3*Δ, *zip1*Δ, or *zip4*Δ fail to form mature tetrads, sporulate, and complete MI (Figures S3A-S3B). These results are consistent with previous reports that the ZMM-dependent class I IH-CO pathway is more important than the Mms4-dependent class II IH-CO pathway in normal meiosis (51,85). Two lines of evidence support that Mms4•Mus81 (and possibly Yen1 and Slx1•Slx4) is subjected to MMR inhibition in hybrid zygotes lacking ZMM protein with anti-MMR activities. First, *P_CLB2_-MSH2* can rescue infertility of hybrid zygotes with *mer3*Δ*, zip1*Δ, *zip4*Δ or *msh4*Δ (Figure 2E). Second, *zip4*Δ *P_CLB2_-MSH2* hybrid zygotes generate much more viable spores than *zip4*Δ *P_CLB2_-MSH2 P_CLB2_-MMS4* and *zip4*Δ *P_CLB2_-MMS4* hybrid zygotes (Figure 5). These findings are not only consistent with a recent report that Msh2 inhibits non-interfering IH-COs in response to genetic polymorphisms in the higher plant *A. thaliana* (169), they also explain (at least in part) why genome-wide mapping of meiotic recombination products in *S. cerevisiae* tetrads with four viable spores revealed that deletion of *MSH2* from SK1/S288c hybrid zygotes only increased COs by ∼20% (4).

### IH-HR products are (normally) produced in *rad51-II3A* and *rad54*Δ hybrid zygotes

To determine if the low fertility phenotype of anti-MMR mutant hybrid zygotes is due to insufficient IH-CO products, we performed physical analyses (Southern hybridization) to quantify both IH-CO and IH-NCO products generated at the *HL* hotspot in the SK1/SK1-*HL* purebred zygotes, as well as those at the *HL\** hotspot and the *HL*-SNP+* hotspot in the SK1/S288c hybrid zygotes, respectively. Genomic DNA was isolated 24 hours after the corresponding diploid zygotes were transferred into sporulation medium, digested by *XhoI* and *NgoMIV*, visualized by Southern hybridization with “probe A”, and then quantified with a TyphoonFLA 9000 biomolecular imager (149). A point mutation was introduced in *HL*\* to mutate the *XhoI* restriction site adjacent to “probe A”. The length of the IH-CO* fragments is longer in both SK1/S288c-*HL*\* and SK1/S288c-*HL*\**-SNP+* hybrid zygotes (∼5.7 kb) than that in the SK1/SK1-*HL* purebred zygotes (∼4.6 kb). The length of the IH-NCO* DNA fragments (∼4.3 kb) is the same in all zygotes (Figures 6A-6D).

**Figure 6.**
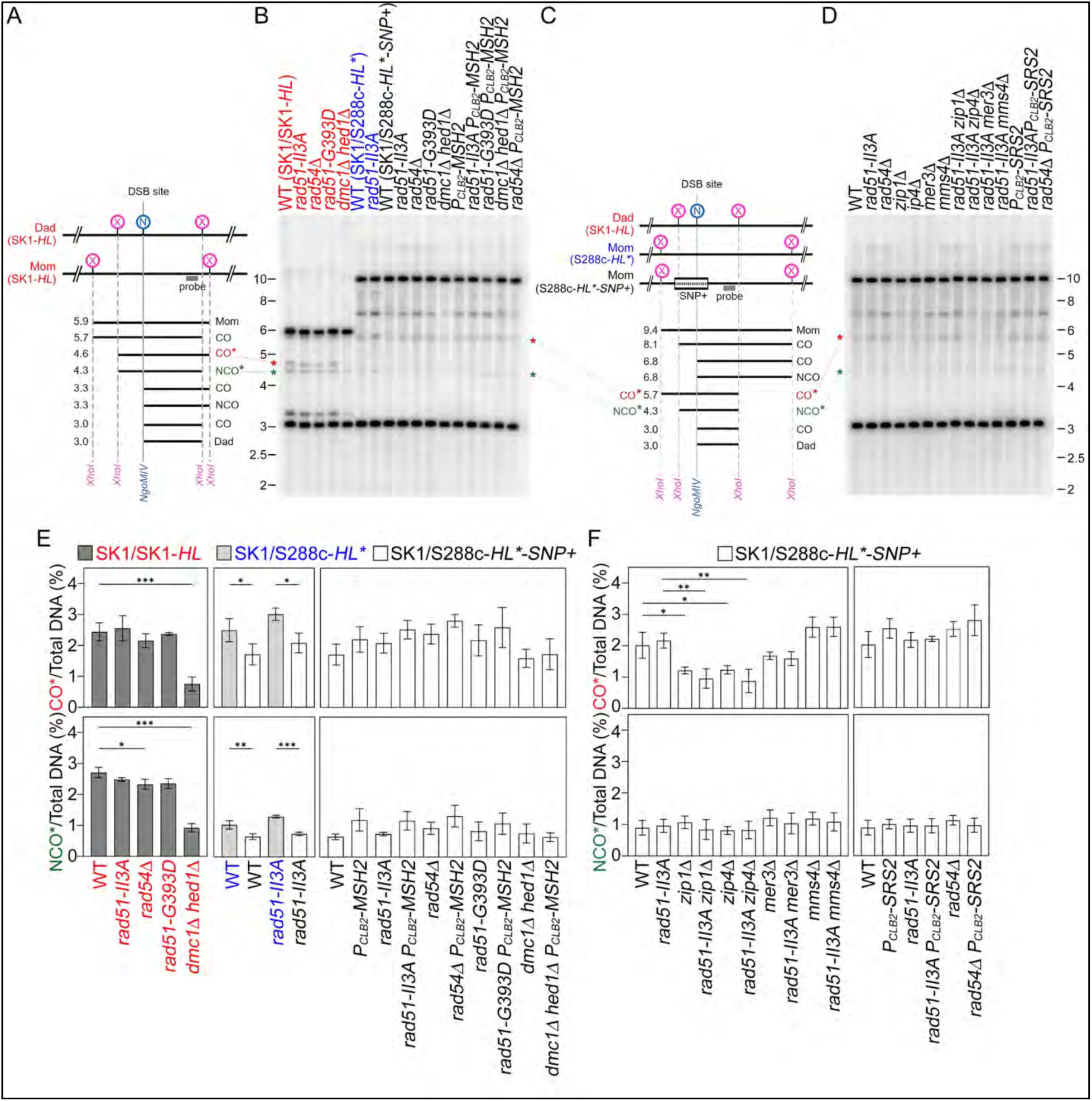
Physical analyses of anti-MMR mutants. (**A,C**) Schematic of the DSB hotspot in SK1/SK1-*HL* purebred zygotes (in red), and in SK1/S288c-*HL\** (in blue) and SK1/S288c-*HL*- SNP+* (in black) hybrid zygotes, respectively. The DNA probe used for Southern hybridization is shown as a grey box. *XhoI* and *NgoMIV* are indicated as X and N, respectively. (**B,D**) Physical analysis by Southern hybridization of recombinants at the *HL* and *HL\** DSB hotspots. Genomic DNA was isolated 24 hours after the meiotic cells were transferred into sporulation medium and then digested with *XhoI* and *NgoMIV* to detect indicated IH-CO* and IH-NCO* recombinants. (**E,F**) Quantification of recombinants from three independent time courses, one of which is shown in B and D. All statistical analyses were carried out as described in Figure 1.

The quantities of both IH-CO* and IH-NCO* products in the SK1/SK1-*HL* purebred zygotes were not affected by *rad51-II3A, rad54*Δ, or *rad51-G393D*, but they were significantly reduced by *dmc1*Δ *hed1*Δ (Figure 6B, lanes 1-5; Figure 6E, left panels). These results are consistent with those of two previous studies on *rad51-II3A* (155) and *dmc1*Δ *hed1*Δ (45). Our DiBAC_4_(5) and DAPI staining experiments also revealed that *dmc1*Δ *hed1*Δ had more profound impacts on sporulation and/or MI than *rad51-II3A*, *rad54*Δ or *rad51-G393D* in both SK1/SK1-*HL* purebred and SK1/S288c-*HL*\**-SNP+* hybrid zygotes, respectively (Figure S2), because the *dmc1*Δ *hed1*Δ meiotic cells accumulated unrepaired DSBs and exhibited persistent DNA damage checkpoint activation and Hop1-T318 phosphorylation (48).

Compared to the SK1/SK1-*HL* purebred zygotes, the increase in genome-wide sequence polymorphism displayed by SK1/S288c-*HL*\* hybrid zygotes had a more profound impact on IH-NCO* than IH-CO* products at this DSB hotspot. The numbers of IH-CO* products in wild-type (*RAD51*) and *rad51-II3A* SK1/SK1-*HL* purebred zygotes (Figure 6B, lanes 1-2; Figure 6E, upper left panel) were similar to those exhibited by wild-type and *rad51-II3A* SK1/S288c-*HL*\* hybrid zygotes (Figure 6B, lanes 6-7; Figure 6E, upper middle panel). In contrast, the quantities of IH-NCO* products in *RAD51* and *rad51-II3A* SK1/SK1-*HL* purebred zygotes (Figure 6B, lanes 1-2; Figure 6E, lower left panel) were significantly higher than those in wild-type and *rad51-II3A* SK1/S288c-*HL*\* hybrid zygotes (Figure 6B, lanes 6-7; Figure 6E, lower middle panel).

The numbers of both IH-CO* and IH-NCO* products in wild-type and *rad51-II3A* SK1/S288c-*HL*\**-SNP+* hybrid zygotes (Figure 6B, lanes 8-9; Figure 6E, middle panels) were lower than those presented by both wild-type and *rad51-II3A* SK1/S288c-*HL*\* hybrid zygotes (Figure 6B, lanes 6-7; Figure 6E, middle panels). Thus, the introduction of 86 exogenous SNPs into the *HL\** hotspot significantly reduced both IH-CO* and IH-NCO* products at this DSB hotspot, supporting our hypothesis that the MMR system can prevent DSB strand exchange between homologous chromosomes in hybrid zygotes (also see below). Notably, the quantities of IH-CO* and IH-NCO* products at this DSB hotspot (Figures 6E and 6F) cannot explicitly explain the low fertility defects of *rad51-II3A* SK1/S288c-*HL*\* and *rad51-II3A* SK1/S288c-*HL*\**-SNP+* hybrid zygotes (Figure 1B and DS2). The same conclusion is applicable to another three anti-MMR mutations of the SK1/S288c-*HL*-SNP+* hybrid zygotes, i.e., *rad54*Δ, *rad51-G393D,* and *dmc1*Δ *hed1*Δ (Figure 6B, lanes 9-12; Figure 6E, right panels). Introduction of *P_CLB2_-MSH2* into wild-type and those four anti-MMR mutants did not significantly alter the quantities of IH-CO* and IH-NCO* products (Figure 6B, lanes 13-17; Figure 6E, right panels).

We also observed that *P_CLB2_-SRS2* does not significantly affect the numbers of IH-CO* and IH-NCO* products at the *HL*-SNP+* hotspot (right panels, Figures 6D and 6F), sporulation efficiency (Figure S3A), and MI progression (Figure S3B) in wild-type, *rad51-II3A*, and *rad54*Δ hybrid zygotes, respectively.

Consequently, we conclude that IH-CO and IH-NCO products both seem to be generated normally by WT and *P_CLB2_-MSH2* hybrid zygotes with *rad51-II3A*, *rad54*Δ, or *P_CLB2_-SRS2*. Therefore, it is unlikely that the hybrid sterility caused by these anti-MMR mutations is due to an insufficiency of IH-HR products at the *HL\** and *HL*\**-SNP+* DSB hotspots.

### *P_CLB2_-MSH2* results in more IH-CO products at the *HL*\**-SNP+* DSB hotspot in *zmm*Δ SK1/S288c-*HL*\*-SNP+ hybrid zygotes

Southern hybridization of wild-type and *rad51-II3A* SK1/S288c-*HL*\*-SNP+ hybrid zygotes revealed that *mer3*Δ, *zip1*Δ and *zip4*Δ do not significantly affect the quantities of IH-NCO* products. In contrast, quantities of IH-CO* products were significantly reduced by *zip1*Δ or *zip4*Δ, slightly reduced by *mer3*Δ, but unaffected by *mms4*Δ (Figures 6D, 6F, and 7A-7D). Thus, both class I IH-CO and IH-NCO products are normally generated in wild-type and *rad51-II3A* hybrid zygotes. We also show that *mer3*Δ*, zip1*Δ, and *zip4*Δ (but not *mms4*Δ) completely prevent sporulation (Figure S3A) and MI progression (Figure S3B) of wild-type and *rad51-II3A* hybrid zygotes. These results are consistent with our hypothesis that Dmc1-mediated IH-HR is more efficient than Rad51-mediated IH-HR in repairing DSBs (32) and inducing SC assembly (94), and that ZMM-dependent class I IH-CO products account for the majority of IH-CO in *S. cerevisiae* (51).

Next, introducing *P_CLB2_-MSH2* into wild-type, *mer3*Δ (Figure 7) and the other four *zmm*Δ mutants (*zip1*Δ, *zip3*Δ, *zip4*Δ and *msh4*Δ; Figures S4A-S4D) increased IH-CO* and/or IH-NCO* products at the *HL*-SNP+* hotspot in the SK1/S288c-*HL*\**-SNP+* hybrid zygotes. The increase of IH-CO* products by *P_CLB2_-MSH2* in *zmm*Δ hybrid zygotes is likely due to the inhibition of Mms4•Mus81 by MMR (Figure 5). *P_CLB2_-MSH2* produced more IH-NCO* products in wild-type and *zmm*Δ hybrid zygotes, showing that Sgs1 can function in an MMR-independent manner, as is the cases in IS-HR and IH-HR between homozygous sequences (86).

**Figure 7.**
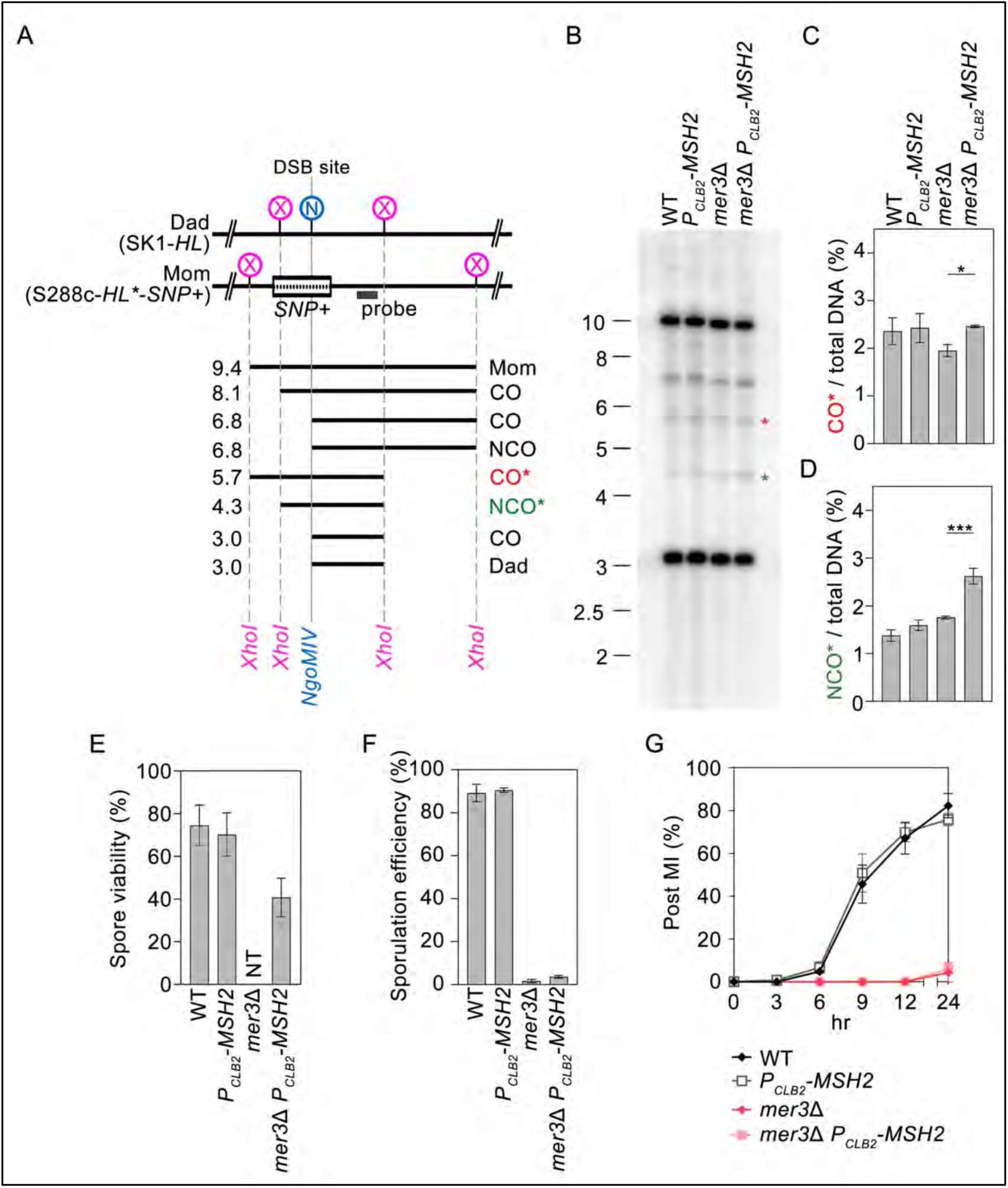
*P_CLB2_-MSH2* increases spore viability and sporulation efficiency but not the quantities of IH-CO and IH-NCO recombinants at the *HL*\* *-SNP+* DSB hotspot of *mer3*Δ SK1/S288c-*HL*\**-SNP+* hybrid zygotes. **(A)** Schematic of the DSB hotspot in the SK1/S288c- *HL*\* *-SNP+* hybrid zygotes. (**B**) Physical analysis by Southern hybridization of recombinants at the *HL* and *HL*\* DSB hotspots. (**C,D**) Quantification of recombinants from three independent time courses, one of which is shown in B. (**E**) Spore viability was determined by tetrad analysis. (**F**) Sporulation efficiency was determined by staining the sporulating cells with DiBAC_4_(5) (141) and analyzed by flow cytometry using a BD FACSymphony A3 Cell Analyzer (143). (**G**) The percentages of cells that completed MI and both MI+MII divisions (Post MI) (139,140) were determined by staining the sporulating cells with DAPI.

DiBAC_4_(5) and DAPI staining experiments revealed that ∼80% wild-type and *P_CLB2_-MSH2* SK1/S288c-*HL*-SNP+* hybrid zygotes generated tetrads and completed MI. Although *P_CLB2_-MSH2* (but not wild-type) SK1/S288c-*HL*-SNP+* hybrid zygotes with *mer3*Δ, *zip1*Δ, or *zip4*Δ could form few mature tetrads that contain viable spores (Figure 7E and Figure S4F), the majority of wild-type and *P_CLB2_-MSH2* SK1/S288c-*HL*-SNP+* hybrid zygotes with *mer3*Δ, *zip1*Δ, or *zip4*Δ sporulated poorly (Figure 7F and S4E) and hardly completed MI (Figure 7G and S4G-S4H). In contrast, more wild-type and *P_CLB2_-MSH2* SK1/S288c-*HL*-SNP+* hybrid zygotes with *msh4*Δ or *zip3*Δ sporulated (Figures S4F), produced viable spores (Figures S4F), and complete MI (Figures S4G-S4H). These results support our hypothesis that Mer3, Zip1 and the ZZS complex possess greater anti-MMR activity than MutSγ and Zip3.

The class I IH-CO pathway accounts for 70-85% of COs in *S. cerevisiae* (51). MMR inhibits Mms4•Mus81 in hybrid zygotes (Figure 5). Three lines of evidence confirm that Mms4•Mus81 functions downstream (but not upstream) of MMR during homeologous recombination: (1) *mms4*Δ does not significantly affect both IH-CO* products and IH-NCO* products in wild-type and *P_CLB2_-MSH2* hybrid zygotes (Figures S5A-S5D). (2) Although the mature tetrads of *mms4*Δ and *mms4*Δ *P_CLB2_-MSH2* hybrid zygotes contain almost no viable spores (S5E), the effects of *mms4*Δ on the capability of wild-type and *P_CLB2_-MSH2* hybrid zygotes to sporulate (Figures S5F) and complete MI (Figures S5G) were similar. (3) *P_CLB2_-MMS4* purebred zygotes generate 72.2±2.8%viable spores, whereas *P_CLB2_-MMS4* and *P_CLB2_-MMS4 P_CLB2_-MSH2* hybrid zygotes generate similar numbers of viable spores (29.2±6.3% and 20.8±7.3%; respectively) (Figure 2C).

### Genome-wide quantities of IH-CO and IH-NCO products in wild-type and *rad51-II3A* SK1/S288c-*HL-SNP+* tetrads with four viable spores

Both the number and position of chromosome-wide IH-COs are tightly controlled by the “CO assurance rule” and the “CO interference rule” (6,57,170,171). To further examine how *rad51-II3A* results in hybrid sterility, we applied Oxford Nanopore third-generation sequencing (TGS) technology and a home-made bioinformatics pipeline TSETA (TGS to Enable Tetrad Analysis) to compare genome-wide IH-CO and IH-NCO profiles in wild-type and mutant SK1/S288c-*HL*\**- SNP+* hybrid zygotes. TSETA is a versatile tool for genome-wide comparison, mapping and visualization of SNPs, InDels, illegitimate mutations, and IH-HR products before and after single meiotic events. The TSETA viewer allows users to instantaneously and continuously visualize all 16 near-complete chromosomes at scales ranging from individual nucleotides to full-length chromosomes (5,132,133,146,147,149).

In brief, four viable spores were isolated by microdissection from the corresponding hybrid zygotes. After germination and vegetative growth, high-quality genomic DNA was purified from the corresponding progeny and subjected to Oxford Nanopore long-read sequencing. As the recommended sequencing depth to generate a high-quality near-complete genome from long-read data varies depending on sequencing type, with 30–50× coverage recommended for Nanopore (172), the sequencing outputs of all haploid strains in this study are 52.7–139.9x (DS3). Next, we applied the “tetrad” mode of TSETA to align and compare the near-complete genome sequences of the two parental haploid strains and the four F1 progeny haploid strains germinated from the corresponding viable spores. We found that the four wild-type tetrads and the four *rad51-II3A* SK1/S288c-*HL*\**-SNP+* tetrads generated 67.0±3.5 and 57.8±6.2 IH-CO products and 47.5±10.0 and 49.5±6.5 IH-NCO products, respectively (DS22).

It was reported previously that one potential caveat with genome-wide sequencing analysis is that IH-CO values may be overestimated due to selection bias because spore viability is reduced in the mutants and only tetrads in which all four spores are viable are used (45). For example, wild-type SK1/S288 hybrid zygotes generated more viable spores than *dmc1*Δ *hed1*Δ and *dmc1*Δ *hed1- 3A* hybrid zygotes (45). *Hed1-3A* encodes a phosphorylation-defective Hed1 mutant, in which the Thr40, Thr41 and Ser42 residues are all substituted with alanine (A) so that the protein cannot be phosphorylated by Mek1 (45). That previous study deployed next-generation sequencing (NGS) using an Illumina HiSeq 2500 instrument to perform genome-wide mapping (with paired-end reads of 150 X 150 bp) of IH-CO and IH-NCO products in 20 wild-type, 12 *dmc1*Δ *hed1*Δ and 12 *dmc1*Δ *hed1-3A* tetrads obtained from SK1/S288c hybrid zygotes. The results revealed little or no effect on the total number of IH-COs between these three types of hybrid zygotes, though IH-NCO events were significantly reduced in *dmc1*Δ *hed1*Δ (45). Previously, we subjected the same NGS reads (45) to TSETA to assemble the corresponding near-complete haploid genome sequences before and after single meiotic events, as well as for genome-wide mapping of IH-CO and IH-NCO products in the 20 wild-type and 12 *dmc1*Δ *hed1*Δ tetrads obtained from the SK1/S288c hybrid zygotes (5). We found that the wild-type and *dmc1*Δ *hed1*Δ tetrads generated similar average numbers of IH-CO and IH-NCO products (5). Moreover, both the median length of GC tracts (∼1000 bp) associated with all IH-HR products and the median density of allelic point disparities (APDs, representing the sum of SNPs and InDels) per CO product (∼4.8 APDs per 1,000 bp) in both wild-type and *dmc1*Δ *hed1*Δ hybrid zygotes proved to be nearly identical (5).

In conclusion, the results of these genome-wide sequencing experiments, like those from physical analysis at the *HL*-SNP+* hotspot, cannot explicitly explain why *rad51-II3A* and *dmc1*Δ *hed1*Δ cause hybrid sterility.

### Quantification of chromosome segregation, CO frequency and interference using linked spore-autonomous fluorescent markers

For meiotic cells unable to form tetrads or produce fertile gametes or viable spores, it is not possible to accurately quantify chromosome-wide IH-HR products using current experimental methods, such as classical genetics, whole-genome sequencing (see above), or immunocytology. In the latter scenario, several “CO designation” protein markers (e.g., *S. cerevisiae* Zip3 and Mlh3, *Caenorhabditis elegans* COSA-1, mouse and *Arabidopsis thaliana* MLH1) can be detected as fluorescent foci along a chromosome under microscopy (76,92,173–175). Although these immunocytological methods do not require genetic markers and can be used in mutants that do not complete meiosis and/or sporulation, concerns have been raised about differences in immunocytological and genetic interferences. For example, several *S. cerevisiae* mutants (e.g., *zip1*Δ, *sgs1*Δ, and *msh4*Δ) show cytological interference but no genetic interference (77).

We overcame the problem of low spore viability by using a spore-autonomous fluorescence system with three pairs of heterozygous alleles on chromosome VIII (138). The purebred zygotes were generated by sexually crossing SK1(*a, VIII, rgC*) with SK1(*α, VIII, RGc*) (138). The fluorescent protein markers in the resulting zygotes allow direct visualization of inheritance patterns in intact yeast tetrads, eliminating the need for microdissection and permitting meiotic segregation patterns to be ascertained even in aneuploid spores. The length of the *CEN8-ARG4- THR1* interval (∼54.9 kb) on chromosome VIII is much longer than that of the *HIS4*::*LEU2-URA3* interval (∼12.0 kb) on chromosome III (61), so it is more suitable for accurately quantifying chromosome-wide CO frequency and CO interference.

To study hybrid meiosis, we constructed intraspecies hybrid zygotes by sexually crossing S288c(*a, VIII, rgC*) with SK1(*α, VIII, RGc*). The *CEN8*-*ARG4* (34,040 bp) and *ARG*-*THR1* (17,684 bp) intervals in the SK1 haploid genome harbor 1.49% (506 bp) and 1.09% (192 bp) SNPs, as well as 2.31% (787 bp) and 2.12% (375 bp) InDels, respectively, relative to those in the S288c haploid genome (DS8). Both the SK1(*a, VIII, rgC*)/SK1(*α, VIII, RGc*) purebred zygotes and the SK1(*α, VIII, RGc*)/S288c(*a, VIII, rgC*) hybrid zygotes produced >80% viable spores (Figure 8A). We confirmed that the spore viability of the SK1/SK1-*HL* purebred zygotes and the SK1/S288c- *HL*-SNP+* hybrid zygotes (Figures 1 and 2, and DS20) is consistent with those obtained from SK1(*a, VIII, rgC*)/SK1(*α, VIII, RGc*) and S288c(*a, VIII, rgC*)/SK1(*α, VIII, RGc*) zygotes (Figure 8A, Table S2, and DS20).

**Figure 8.**
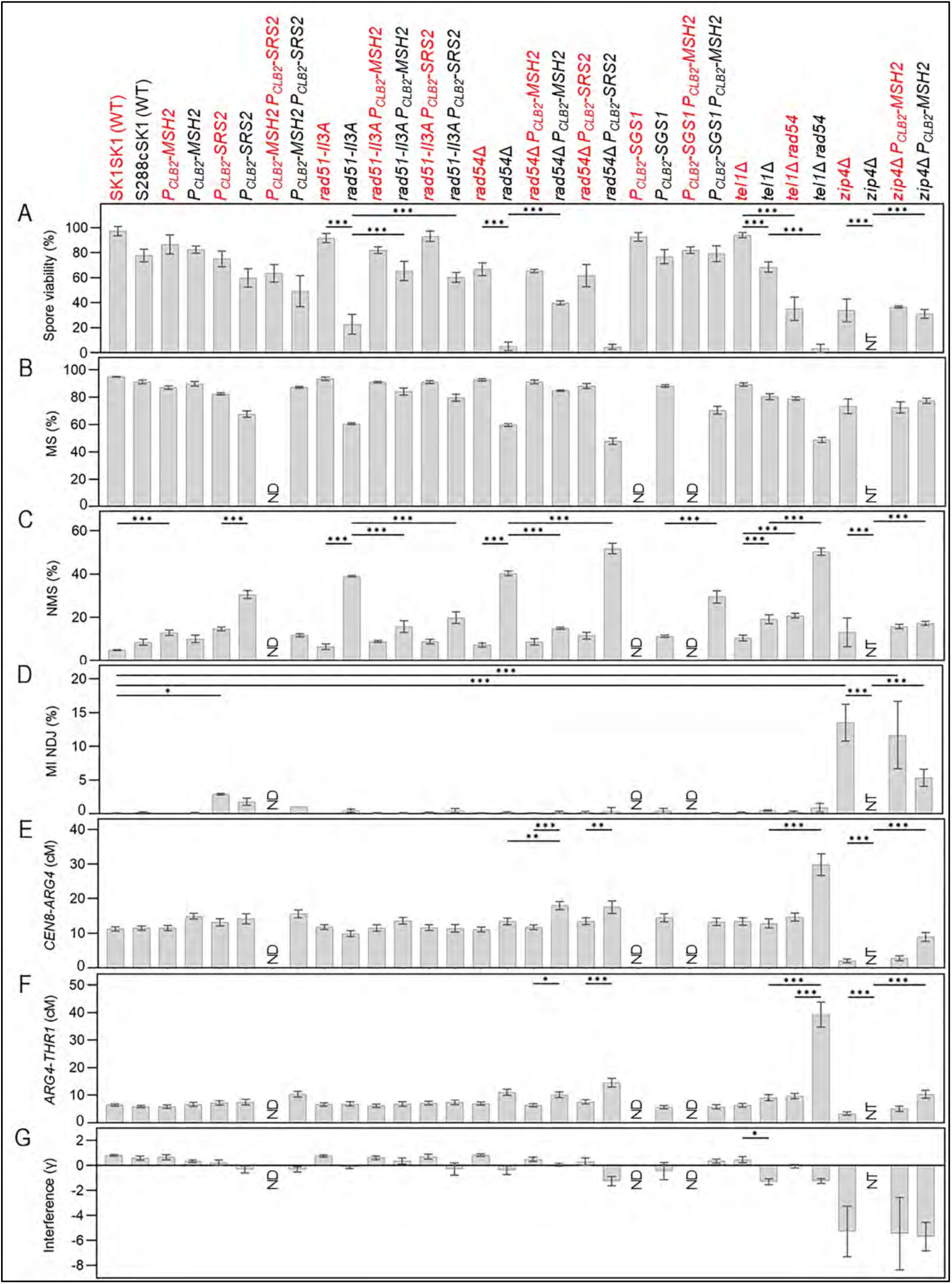
Spore viability and chromosomal segregation patterns of the indicated SK1(*a, VIII, rgC*)/SK1(*α, VIII, RGc*) purebred zygotes (in red) and SK1(*α, VIII, RGc*)/S288c(*a, VIII, rgC*) hybrid zygotes (in black). **(A)** Spore viability of indicated purebred and hybrid zygotes. (**B**) The percentages of tetrads with all three heterozygous fluorescent marker genes that underwent Mendelian segregation (MS) or 2^+^:2^-^ segregation. (**C**) The percentages of tetrads with one of the three fluorescent marker genes that underwent non-Mendelian segregation (NMS), including 3^+^:1^−^, 1^+^:3^−^, 4^+^:0^−^, and 0^+^:4^−^ segregation. (**D**) The percentages of tetrads with the three fluorescent marker genes that underwent MI nondisjunction (NDJ). (**E**) Genetic distances between *CEN8* and *ARG4*. (**F**) Genetic distances between *ARG4* and *THR1*. (**G**) CO interference between the *CEN8*-*ARG4* and *ARG4*-*THR1* intervals. NT (no or very few tetrads). ND (not determined).

Tel1^ATM^ is known to regulate the number, local clustering, and genome-wide location of meiotic DSBs (42,76,176,177). We found that the order of spore viability is WT SK1/SK1 ∼ *tel1*Δ SK1/SK1 > WT S288c/SK1 > *tel1*Δ S288c/SK1 ∼ *rad54*Δ SK1/SK1 > *rad54*Δ *tel1*Δ SK1/SK1 >> *rad54*Δ S288c/SK1 ∼ *rad54*Δ *tel1*Δ S288c/SK1 (Figure 8A). These results indicate that although *tel1*Δ reduces the fertility of both purebred and hybrid zygotes, Rad54’s anti-MMR activity is preferentially required for normal meiosis in wild-type and *tel1*Δ hybrid zygotes than in wild-type and *tel1*Δ purebred zygotes. The reduced hybrid fertility caused by *tel1*Δ could be due to the fact that loss of *TEL1* induces excessive and clustered DSBs, leading to either more residual DSBs and continuous DNA damage checkpoint activation (as also the case for *dmc1*Δ *hed1*) in Phase II (48) or more closely spaced IH-CO products (see below).

Temperature is a key factor regulating meiosis in *S. cerevisiae* and other sexual eukaryotes (e.g., wheat). For example, *S. cerevisiae zmm*Δ mutants (but not wild-type) undergo sporulation better at lower temperate (23 °C) than at higher temperatures (30 °C and 33 °C) (51). Dmc1 is a candidate protein endowing temperature tolerance during wheat meiosis (178), with the wheat *DMC1-D1* allele being responsible for better preserving normal CO formation at low temperatures relative to at high temperatures (179). Here, we report that temperature preferentially affects the anti-MMR activities of the ZZS complex, but not those of Rad51 and Rad54. First, like WT purebred and hybrid zygotes, those with *rad51-II3A*, *rad54*Δ, *rad51-II3A P_CLB2_-MSH2*, or *rad54*Δ *P_CLB2_-MSH2* generated similar levels of viable spores at 23 °C and 30 °C, respectively (Figure 9). Second, the order of spore viability of purebred zygotes with *zip4*Δ and *zip4*Δ *P_CLB2_-MSH2* is 30 °C > 23 °C ∼ 33 °C. In contrast, hybrid zygotes with *zip4*Δ and *zip4*Δ *P_CLB2_-MSH2* produced 11% and 16% viable spores at 23 °C, but hardly formed mature tetrads at 30 °C and 33 °C. *P_CLB2_-MSH2* increased *zip4*Δ hybrid fertility at 30 °C and 33 °C, i.e., 31% and 23% viable spores, respectively (Figure 9). Accordingly, in *zip4*Δ hybrid zygotes, high temperatures may promote more stable binding between mismatched base pairs and the MMR system than low temperatures. Based on our results in Figure 9 and those from studies on wheat meiosis (178,179), we propose that high temperature preferentially affects the Dmc1-mediated IH-HR pathway relative to the Rad51-mediated IH-HR pathway in hybrid zygotes displaying high sequence heterogeneity. This hypothesis is consistent with the fact that Dmc1-made joint molecules can effectively induce complete SC formation in Phase I, whereas the SC induced by Rad51-generated joint molecules in *dmc1*Δ *hed1*Δ meiotic cells is inefficient and abnormal (94).

**Figure 9.**
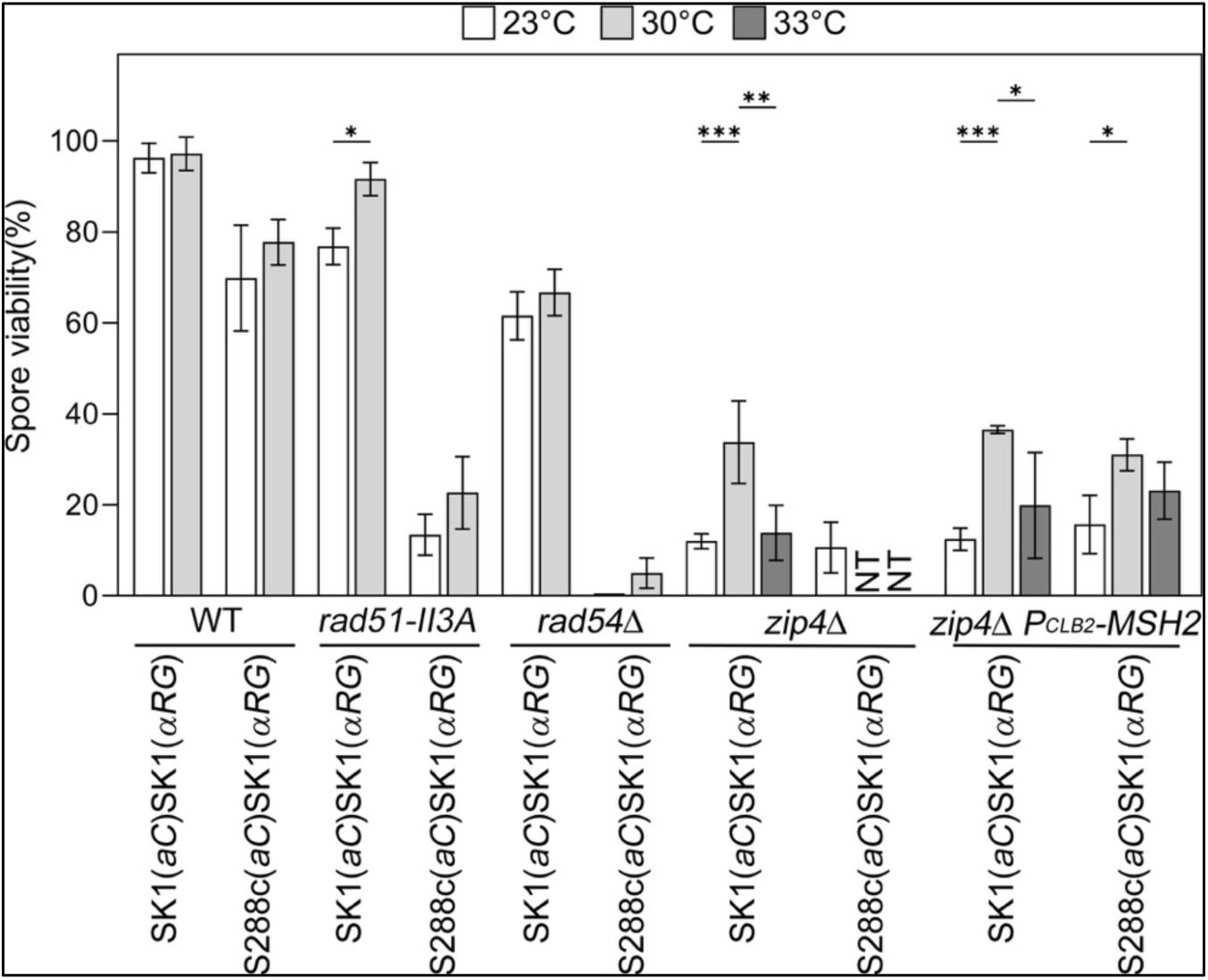
Spore viability of the indicated SK1(*a, VIII, rgC*)/SK1(*α, VIII, RGc*) purebred zygotes (in red) and SK1(*α, VIII, RGc*)/S288c(*a, VIII, rgC*) hybrid zygotes at 23 °C (white), 30 °C (grey), or 33 °C (black).

Next, we quantified the percentages of tetrads exhibiting Mendelian segregation (MS; Figure 8B), non-Mendelian segregation (NMS; Figure 8B), and MI NDJ (Figure 8C) at 30 °C. Since allelic conversion (IH-NCO) occurs between heterozygotic sites and results in mismatched base pairs that are then recognized and corrected by the MMR system, one of the alleles is converted to the other. This can lead to NMS, i.e., 3^+^:1^−^, 1^+^:3^−^, 4^+^:0^−^, or 0^+^:4^−^ alleles in tetrads (based on the four double-stranded DNA duplexes present). The percentages of tetrads that underwent 2^+^:2^−^ 3^+^:1^−^, 1^+^:3^−^, 4^+^:0^−^, or 0^+^:4^−^ segregation from each pair of heterozygous spore-autonomous fluorescent marker genes were determined, respectively (Figure S6). The NMS tetrads include all tetrads with ≥1 non-2^+^:2^−^ heterozygous spore-autonomous fluorescent marker genes (Figure 8C). The MI NDJ tetrads present two white spores and two black spores under fluorescence microscopy (Figure 8D). Although this segregation pattern might also arise from a four-strand double crossing over (DCO) event in the *CEN8-ARG4-THR1* interval, it has been reported previously that the frequency of four-strand DCOs is significantly lower than that of MI NDJ (138). The MS tetrads harbor three 2^+^:2^−^ fluorescent protein markers (Figure 8B) and were used to quantify genetic distances (Figures 8E and 8F) and CO interference [γ = 1 – coefficient of coincidence (c.o.c.)] (Figure 8G). Genetic distances were calculated using Perkins equation: cM (centimorgan) = [100 (6NPD + T)] / [2 (PD + NPD + T)], where PD, NPD and T are parental ditype, non-parental ditype, and tetrad type, respectively. Both “genetic distances ± standard errors” and the “NPD ratio” were determined using Stahl Laboratory Online Tools (https://elizabethhousworth.com/StahlLabOnlineTools) (150). The NPD ratio (*fN_obs_*/*fN_exp_*) is an indication of the intensity of interference between two neighboring genetic markers (i.e., *CEN8* and *ARG4* or *ARG4* and *THR1*), with smaller values indicating stronger positive interference (151,152) (Table S2 and DS23). The c.o.c. values were calculated by dividing the actual frequency of double recombinants between the *CEN8-ARG4* and *ARG4-THR1* intervals by the expected frequency (74). A positive γ value indicates that the spacing of IH-CO events departs from a random distribution, whereas a negative γ value reflects IH-CO events that are more clustered than the null expectation. We present the outcomes of these analyses for the different zygote types below.

(1) NMS

The wild-type purebred and hybrid zygotes produced ∼5% and ∼13% NMS tetrads, whereas *P_CLB2_- MSH2* purebred and hybrid zygotes generated ∼13% and ∼10% NMS tetrads, respectively. *P_CLB2_-* Srs2 and Sgs1 respond differently to the MMR system during homeologous recombination. *P_CLB2_- SRS2* hybrid zygotes produced ∼30% NMS tetrads. *P_CLB2_-SGS1* hybrid zygotes generating approximately 11% NMS tetrads. Hybrid zygotes with *P_CLB2_-SRS2 P_CLB2_-MSH2* and *P_CLB2_-SGS1 P_CLB2_-MSH2* had approximately 12% and 29% NMS tetrads, respectively (Figure 8B and DS23).

*rad51-II3A*, *rad54*Δ, and *tel1*Δ induced more allelic conversion in hybrid zygotes than in purebred zygotes. Both *P_CLB2_-MSH2* and *P_CLB2_-SRS2* reduced the percentages of NMS tetrads generated by the *rad51-II3A* hybrid zygotes, i.e., from 39% to 16% and 20%, respectively. In contrast, the order of NMS tetrad percentages in three *rad54*Δ hybrid zygotes is *P_CLB2_-MSH2 rad54*Δ (∼15%) < *rad54*Δ (∼40%) < *P_CLB2_-SRS2 rad54*Δ (∼52%). The order of NMS tetrad percentages in *tel1*Δ and *tel1*Δ *rad54*Δ zygotes is *tel1*Δ SK1/SK1 (∼%11) < *tel1*Δ *rad54*Δ *P_CLB2_- MSH2* S288c/SK1 (∼19%) ∼ *tel1*Δ *rad54*Δ SK1/SK1 (∼21%) < *tel1*Δ *rad54*Δ S288c/SK1 (∼50%). These results are consistent with our hypotheses. First, that Srs2 antagonizes Rad51’s anti-MMR activity by dissembling Rad51-ssDNA nucleoprotein filaments. Second, that Rad54 acts after Srs2 during Rad51-mediated D-loop formation. Third, although *tel1*Δ induces excessive and clustered DSBs during meiosis, Rad54’s anti-MMR activity outcompetes its pro-HR function in both WT and *tel1*Δ hybrid zygotes.

The ZZS complex promotes class I IH-CO formation and also antagonizes MMR in hybrid zygotes. Accordingly, in *zip4*Δ hybrid zygotes, MMR inhibits class II IH-CO formation by Mms4•Mus81, Yen1 and Slx1•Slx4, and it promotes IH-NCO formation (allelic conversion) by Sgs1, potentially explaining why *zip4*Δ hybrid zygotes hardly form mature tetrads (Figure 8A). In contrast, *zip4*Δ *P_CLB2_-MSH2* hybrid zygotes, *zip4*Δ purebred zygotes, and *zip4*Δ *P_CLB2_-MSH2* purebred zygotes generated mature tetrads with 31-36% viable spores (Figure 8A), including ∼17%, ∼13%, and ∼16% NMS tetrads, respectively (Figure 8C). These results are in agreement with our suggestion that Sgs1 can be activated in an MMR-independent manner to form IH-NCO products and that ZMM proteins can inhibit MMR-dependent and -independent functions of Sgs1.

(2) MI NDJ

Most of the purebred and hybrid zygotes we examined in Figure 8 generated almost no MI NDJ tetrads (<1%). *P_CLB2_-SRS2* induced about ∼4% and ∼2% MI NDJ tetrads in purebred and hybrid zygotes, respectively. *zip4*Δ induced ∼13%, ∼12% and ∼5% MI NDJ tetrads in wild-type, *P_CLB2_- MSH2* purebred, and *P_CLB2_-MSH2* hybrid zygotes, respectively (Figure 8D). Moreover, *zip4*Δ hybrid zygotes produced almost no mature tetrads (Figure 8A), probably because of a much higher frequency of chromosome missegregation.

(3) Genetic distances and CO interference

Wild-type purebred and hybrid zygotes presented similar CO frequencies in the *CEN8-ARG4* interval (11.23 cM and 11.44 cM) and the *ARG4-THR1* interval (6.41cM and 6.81 cM), and similar positive CO interferences between the *CEN8-ARG4* and *ARG4-THR1* intervals (γ = 0.82 and 0.60, respectively) (Figures 8E-8G).

The ZZS complex promotes formation of class I IH-COs in purebred zygotes (51). Indeed, *zip4*Δ and *zip4*Δ *P_CLB2_-MSH2* purebred zygotes exhibited reduced CO frequencies in the *CEN8- ARG4* interval (1.96 cM and 2.74 cM) and the *ARG4-THR1* interval (3.33 cM and 5.06 cM). Both class I and class II IH-CO formation are diminished in *zip4*Δ hybrid zygotes because the MMR system not only inhibits class II IH-CO formation by Mms4•Mus81, but also promotes IH-NCO formation by Sgs1 in *zip4*Δ hybrid zygotes. Therefore, *zip4*Δ hybrid zygotes produce almost no mature tetrads, probably because of much lower CO frequencies and a higher NMS frequency. *P_CLB2_-MSH2* significantly increased the CO frequencies of *zip4*Δ hybrid zygotes (8.90 cM and 10.31 cM). Notably, *zip4*Δ *P_CLB2_-MSH2* hybrid zygotes, *zip4*Δ purebred zygotes, and *zip4*Δ *P_CLB2_- MSH2* purebred zygotes generated similar amounts of viable spores (31.0%, 33.8% and 36.6%; Figure 8A, Table S2 and DS23) and exhibit similar negative CO interferences (γ= -5.68, -5.27, and -5.54; Figure 8G, Table S2 and DS23). Therefore, neither the total amount nor the non-interfering nature of IH-COs in these *zip4*Δ zygotes significantly affects their fertility.

Most of the non-*zmm*Δ purebred and hybrid zygotes we examined in Figure 8 display CO frequencies similar to WT. *tel1*Δ *rad54*Δ induced the highest CO frequencies in both purebred and hybrid zygotes in the *CEN8-ARG4* interval (14.66 cM and 29.83 cM) and the *ARG4-THR1* interval (9.69 cM and 39.29 cM), respectively. *P_CLB2_-SRS2 P_CLB2_-MSH2* (15.57 cM and 10.32 cM)*, rad54*Δ *P_CLB2_-MSH2* (17.98 cM and 10.16 cM), and *rad54*Δ *P_CLB2_-SRS2* (17.52 cM and 14.55 cM) also significantly increased CO frequencies in hybrid zygotes. Apart from the *tel1*Δ *rad54*Δ purebred zygotes (γ= -0.05), all other non-*zmm*Δ purebred zygotes exhibited positive CO interference (γ= 0.25 to 0.77). Five non-*zmm*Δ hybrid zygotes exhibited positive CO interference (γ= 0.25 to 0.77), including *P_CLB2_-MSH2*, *rad51-II3A P_CLB2_-MSH2*, *rad54*Δ *P_CLB2_-MSH2*, and *P_CLB2_-MSH2 P_CLB2_- SGS1.* In contrast, the other nine non-*zmm*Δ hybrid zygotes exhibited negative CO interference (γ= -0.13 to -1.3), including *P_CLB2_-SRS2, P_CLB2_-SRS2 P_CLB2_-MSH2, rad51-II3A, rad51-II3A P_CLB2_- SRS2, rad54*Δ, *rad54*Δ *P_CLB2_-SRS2, P_CLB2_-SGS1, tel1*Δ, and *tel1*Δ *rad54*Δ.

## Discussion

We propose a hypothetical model (Figure 10) to explain how Rad51 and Dmc1 exhibit similar mismatch tolerability during homology-directed DSB repair *in vivo* (159). In Phase I of meiosis, Rad51 is inhibited by Hed1. Dmc1 tolerates mismatched base pairs and catalyzes D-loop formation. The mismatched base pairs in the Dmc1-generated D-loops are then recognized by Msh2- containing complexes, thereby facilitating Sgs1-mediated IH-NCO formation and preventing Mms4•Mus81 (and probably Yen1 and Slx1•Slx4) from forming class II IH-COs during early meiosis. This hypothesis is not only consistent the report that Msh2 inhibits non-interfering IH-COs in response to genetic polymorphisms in the higher plant *A. thaliana* (169), but also explains (at least in part) why genome-wide mapping of meiotic recombination products in *S. cerevisiae* tetrads with four viable spores revealed that deletion of *MSH2* from SK1/S288c hybrid zygotes only increased COs by ∼20% (4). Subsequently, ZMM proteins (e.g., Mer3) are recruited to Dmc1- made D-loops and/or the resulting joint molecules to antagonize Sgs1 and MMR, thereby promoting the formation of class I and class II IH-COs and limiting allelic conversion (IH-NCOs) by Sgs1. Dmc1 is a potent inhibitor of Srs2. Accordingly, the infertility phenotypes of *zip4*Δ hybrid zygotes can be rescued by *P_CLB2_-MSH2*, *P_CLB2_-SGS1*, *P_CLB2_-MSH2 P_CLB2_-SGS1*, and *P_CLB2_-MSH2 P_CLB2_-SRS2*, but not by *P_CLB2_-SRS2*, *P_CLB2_-MMS4* or even *P_CLB2_-MSH2 P_CLB2_-MMS4*. Our results also indicate that Sgs1 can function in an MMR-independent manner during IS-HR or IH-HR between the same maternal and paternal genomic sequences. This MMR-independent function of Sgs1 is antagonized by the ZMM proteins that mediate SC assembly (86). Notably, IH-NCO can be generated in a Sgs1-independent manner because *P_CLB2_-SGS1* and *P_CLB2_-SGS1 P_CLB2_-MSH2* hybrid zygotes generated approximately 12% and 29% NMS tetrads, respectively (Figure 8C). It will be interesting to investigate whether other HR proteins are responsible for IH-NCO formation in *P_CLB2_-SGS1* and *P_CLB2_-SGS1 P_CLB2_-MSH2* hybrid zygotes.

**Figure 10.**
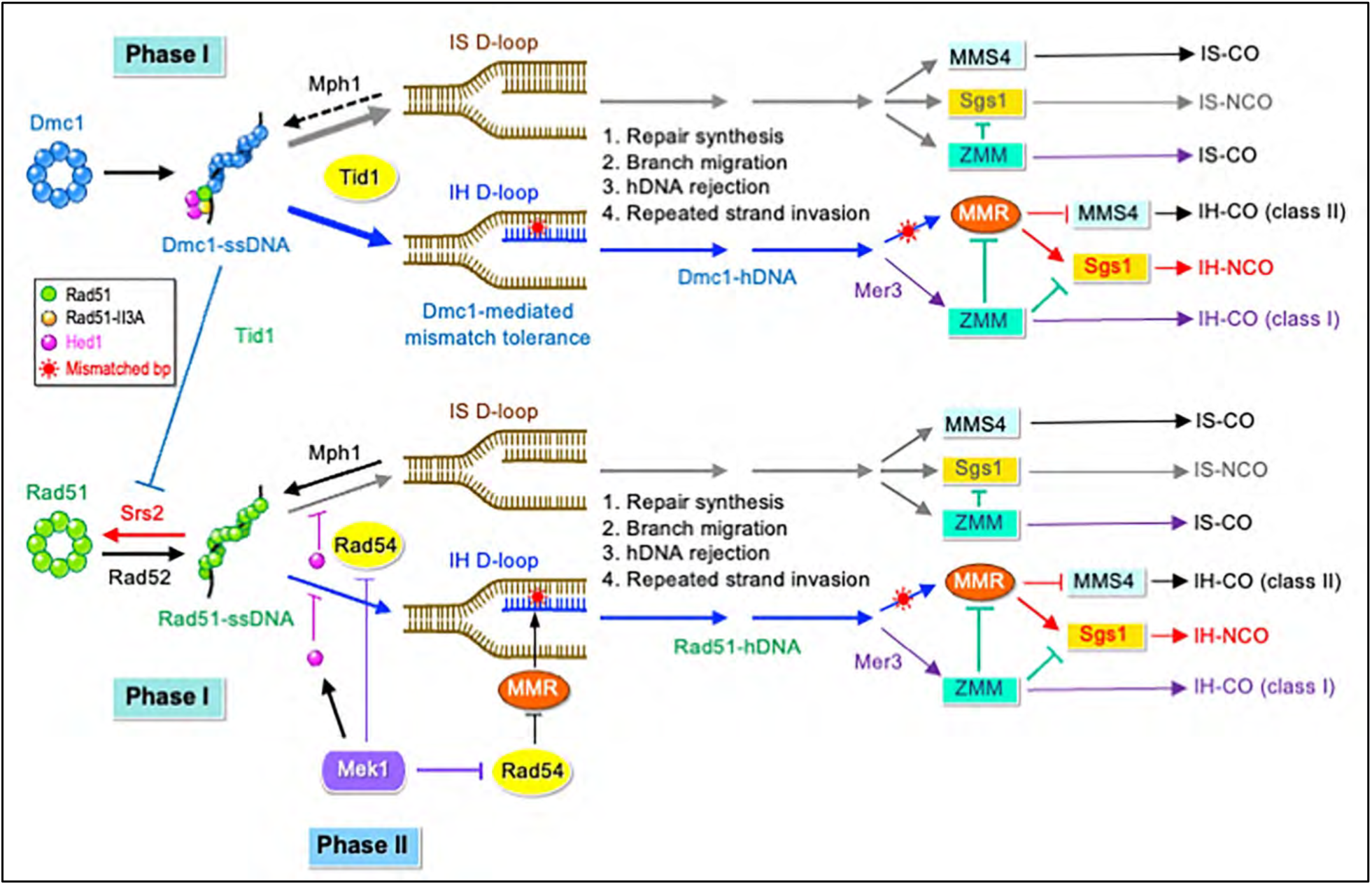
Roles of anti-MMR proteins in Dmc1- and Rad51-mediated homology-directed DSB repair during Phase I and Phase II of intraspecies hybrid meiosis. IH: interhomolog; IS: intersister. _*_: mismatched base pair (bp; in red).

In Phase II of meiosis, residual DSBs activate the DNA damage checkpoint and inactivate Hed1, thereby activating Rad51 (45). Rad51’s accessory proteins (e.g., Rad52, Rad59, and Rad55•Rad57) compete with Srs2 to promote the formation of Rad51-ssDNA presynaptic filaments. These latter then recruit Rad54 to facilitate the formation of Rad51-made D-loops. During D-loop formation, the anti-MMR activity of Rad51 prevents mismatched base pairs from being recognized and corrected by three MMR protein complexes (MutSα, MutSα, and MutLα). After the formation of Rad51-made D-loops and/or during the generation of resulting joint molecules, Rad54’s anti-MMR activity acts through its chromatin remodeling and/or branch migration functions to exclude the Msh2-containing MMR complexes. Since both Dmc1 and Hed1 are absent from *dmc1*Δ *hed1*Δ hybrid zygotes, many DSBs are not properly repaired, leading to continuous activation of the DNA damage checkpoint and Rad54 inhibition via Mek1-mediated phosphorylation. Rad54’s anti-MMR activity becomes indispensable to facilitate Rad51-mediated IH-HR in hybrid zygotes. Accordingly, both *P_CLB2_-MSH2* and *P_CLB2_-SRS2* can rescue the hybrid infertility induced by *rad51-II3A.* In contrast, only *P_CLB2_-MSH2*, but not *P_CLB2_-SRS2*, can rescue the hybrid infertility induced by *rad54*Δ*. tel1*Δ induces an excess of and clustering of DSBs, leading to many residual DSBs in Phase II of meiosis, as in the case of *dmc1*Δ *hed1*Δ, explaining why Rad54’s anti-MMR activity is also more important in *tel1*Δ hybrid zygotes than in *tel1*Δ purebred zygotes.

The most important discovery of this study is that the SC evolved to counter the MMR system in response to sequence heterogeneity. Based on our hypothetical model presented in Figure 10, formation of both class I and class II IH-COs is diminished in *zip4*Δ hybrid zygotes. In contrast, only class I IH-CO formation, but not class II IH-CO formation, is blocked in *zip4*Δ purebred zygotes. These scenarios explain why *zip4*Δ hybrid zygotes exhibit much more severe sterility phenotypes than *zip4*Δ purebred zygotes, as well as at higher temperatures (30 °C and 33 °C) than at a lower temperature (23 °C). It has been reported previously that MutLα forms a dynamic ensemble with a meiosis-specific DNA helicase, Mer3, at Dmc1-dependent D-loops to limit Sgs1- dependent NCO formation *in vivo* (64,65). Accordingly, it will be interesting to investigate further if Mer3 and other ZMM proteins possess different antagonistic preferences toward MMR components in *S. cerevisiae*, including MutSα, MutSα, MutLα, MutLα, and MutLγ.

The SC exists in many (but not all) sexual reproductive eukaryotes, so it will also be important to determine if synapsis-promoting ZMM proteins in other sexual eukaryotes also exhibit anti-MMR activities in hybrid zygotes and how species that have lost the SC during their evolution antagonize the MMR system in hybrid zygotes. For example, independent intraspecies hybrid zygotes have been generated by sexually crossing two isogenic *S. pombe* haploid strains that, after moderate chemical mutagenesis, produced high levels (>94%) of viable spores and unusually large numbers of COs per chromosome (180). *S. pombe* does not form a fully mature SC, though it does form cytologically distinct linear elements (LEs) that exhibit similarities to SC-derived axial elements (AE). *S. pombe* LEs and *S. cerevisiae* AEs display common features, and the key structural components in these two organisms, i.e., Rec10 (*S. pombe*) and Red1 (*S. cerevisiae*), exhibit some amino acid conservation (181). Further investigations will reveal if the components of *S. pombe* LEs and/or *S. cerevisiae* AEs exhibit anti-MMR activity.

We also report for the first time that only *S. cerevisiae* Rad51 and Rad54, but not Dmc1 or Tid1, possess anti-MMR activities. *S. cerevisiae* Rad51 and Rad54 are ubiquitously expressed in both mitotic and meiotic cells, whereas Dmc1 and its biological binding partner Tid1/Rdh54 are meiosis-specific. Both Rad51 and Dmc1 exist in many other sexual eukaryotes, including the fission yeast *S. pombe*, basidiomycete fungi, higher plants, and mammals. In contrast, Dmc1 is absent from Rad51-only species, including *Drosophila melanogaster*, *Caenorhabditis elegans*, and some Pezizomycotina filamentous fungi (e.g., *Neurospora crassa*, *Sodaria macrospora*, and *Trichoderma reeesi*). We have reported previously that ancestral *T. reeesi* Rad51 evolved to acquire Dmc1-like properties by creating multiple structural variations (5). Consequently, not only is it essential for IH-HR during both purebred and hybrid meiosis, but it also possesses better mismatch tolerance than *S. cerevisiae* Rad51 during D-loop formation *in vitro* and IH-HR *in vivo* (5). It will be interesting to investigate in future if Rad51 and Dmc1 in other sexual eukaryotes also possess anti-MMR activity to antagonize the MMR system in hybrid zygotes.

Most studied sexual eukaryotes, such as *S. cerevisiae*, possess at least one Rad54 paralog, including *S. pombe* Rad54 and Tid1, human and mouse (Rad54A and Rad54B) (182), *D. melanogaster* RAD54 (also called OKRA or OKA) (183), *C. elegans* RAD-54.L and RAD-54.B (184), *N. crassa* MUS-25 and SAD-6 (185–187), and *T. reesei* RAD54-like protein (5). The amino acid sequences of *S. pombe* Tid1 and Rad54, as well as that of *N. crassa* MUS-25, are more similar to those of *S. cerevisiae* Tid1 than Rad54 (187). *S. pombe* Tid1 and Rad54 cooperate to repair meiotic DSBs (188). *N. crassa* MUS-25 is important for both MEI-3 (Rad51)-mediated DSB repair (185) and a vegetative HR-dependent RNA silencing process called “quelling” (186). SAD-6 is involved in meiotic silencing by unpaired DNA (MSUD), a process that detects unpaired regions between homologous chromosomes and silences them for the duration of sexual development (187). Only Rad54A, but not Rad54B, is needed for a normal distribution of Rad51 on meiotic chromosomes in mice (182). *C. elegans* RAD-54.L is essential for removing RAD-51 from meiotic DSB sites to enable recombination to progress, whereas RAD-54.B is largely dispensable for meiotic DSB repair but required to prevent RAD-51 hyperaccumulation on unbroken DNA during the meiotic sub-stage when DSBs and early recombination intermediates form (184). *D. melanogaster* RAD54*-*deficient flies develop normally, but the females are sterile (189). Further studies will reveal if the Rad54 paralogs in other sexual eukaryotes also exhibit anti-MMR activities in hybrid zygotes.

Meiosis is highly related to speciation. Based upon the findings in our study, we propose that an imbalance between MMR and HR may contribute to the post-zygotic isolation between two closely related species, e.g., *Saccharomyces cerevisiae* W303 and *Saccharomyces paradoxus* N17 (129). Due to a higher degree of sequence heterogeneity in the W303/N17 hybrid zygotes than in SK1/S288c intraspecies hybrid zygotes, amounts of HR proteins with anti-MMR activities in the former may not be sufficient to prevent the MMR system from stably associating with mismatched base pairs in D-loops and/or joint molecules. According to our hypothetical model presented in Figure 10, the W303/N17 hybrid zygotes may produce too many Sgs1-mediated IH-NCO products and insufficient IH-COs, thereby leading to hybrid sterility. This scenario eloquently explains why the fertility of W303/N17 hybrid zygotes can be improved to the level of non-hybrids by introducing *P_CLB2_-SGS1* (107,128,129) and *P_CLB2_-SGS1 P_CLB2_-MSH2* (129). Consequently, we conclude that the hybrid infertility of W303/N17 zygotes is not due to insufficient IH-HR, but rather to excessive IH-NCOs or even abnormal MMR-induced multichromatid joint molecules and IH-COs. Quantification of the IH-NCO and IH-CO products generated in W303/N17 hybrid zygotes using the spore autonomous fluorescent method utilized herein would be an appropriate way of testing this interesting hypothesis.

Although the spore autonomous fluorescent method is superior to several other experimental approaches (e.g., classical genetics, genome-wide sequencing, physical analysis, and cytology) for accurate quantification of genetic distance, CO interference, and allelic conversion (IH-NCO), we acknowledge that it has several potential drawbacks. First, only tetrads (but not dyads and triads) were used herein to quantify genetic distances and CO interference, as it is difficult to quantify IH-HR products in meiotic cells that do not sporulate well (i.e., monads, dyads or triads). Second, we show that intraspecies hybrid zygotes with anti-MMR mutations produce significant amount of NMS tetrads, so further investigations are needed to establish if DCO events that cause negative CO interference are indeed attributable to two authentic IH-CO products.

## Acknowledgments

We thank both Imaging Core Laboratory and Genomic Core Laboratory of the Institute of Molecular Biology, Academia Sinica, for technical support, John O′Brien for English editing, Matthew Neale (University of Sussex, UK) for sharing the PCR-based *P_CLB2-_HA::KanMX4* integration plasmid, Scott Keeney (HHMI and Sloan Kettering Institute, USA) for sharing the SK1 strains and/or integration plasmids with three spore-autonomous fluorescence markers, Valérie Borde for sharing both *zip2-ΔXPF* and *zip4-N919Q* mutants, Eric Alani (Cornell University) for sharing the *sgs1*Δ-C795 mutant, Valérie Borde (Institut Curie, France), Michael Lichten (NIH, USA), Gerry Smith (Fred Hutchinson Cancer Center, USA), and Nancy Kleckner (Harvard University, USA) for their helpful research advice, and Su-Chien Chen, Sheng-Yuan Chen, and Agatha Cecilia Sihite (IMB, Academia Sinica) for research assistance.

## Supplementary data

Supplementary Tables S1 and S2, figures S1-S6, and Supplemental datasets DS1-DS22

## Data availability

All relevant data are within the paper and its Supporting information files. The bioproject accession number of Nanopore sequencing datasets is PRJNA1216236.

## Consent for publication

All authors have read and approved of its submission to this journal.

## Supplementary materials

**Supplemental Figures S1-S6 and Tables S1-S2 are provided in three separate PDF files, respectively.**

**Datasets DS1-DS16.** List of SNPs and InDels between 16 individual chromosomes of the haploid genomes of SK1 and S288c, respectively. The strain numbers of the SK1 and S288c haploid strains are WHY13008 and WHY14835, respectively. All SNPs and InDels within the two telomeres of all 16 chromosomes (in black), the rDNA loci on the tenth chromosome (in blue), and the *CEN8- ARG4-THR1* intervals on the eighth chromosome (in red) are indicated.

**Dataset DS17. Numbers and percentages of SNPs and InDels between the near-complete haploid genomes of SK1 and S288c.**

**Dataset DS18. List of yeast strains, numbers, and genotypes.**

**Dataset DS19. Nucleotide sequences of the four chromosomal regions shown in Figure 1A.**

**Dataset DS20. Raw data on spore viability.** All diploid cells were incubated at 30 °C in sporulation medium for 48 hours and then scored for their capability to form mature tetrads. Spore viability was determined by isolating the four single spores from each tetrad on YPDA plates.

**Dataset DS21. Uncropped immunoblots of Figure S1.**

Dataset DS22. Genome-wide mapping of interhomolog CO and NCO products generated by four wild-type and four *rad51-II3A* SK1/S288C-*HL*\**-SNP+* hybrid zygotes. The genomic DNA of four representative F1 progeny was determined by PacBio SMRT technology and analyzed by TSETA to identify genome-wide meiotic recombination events, as described previously (5,132,133).

**Dataset DS23. Raw spore-autonomous fluorescence datasets presented in Figure 8 and Table S2.**

**Figure S1.**
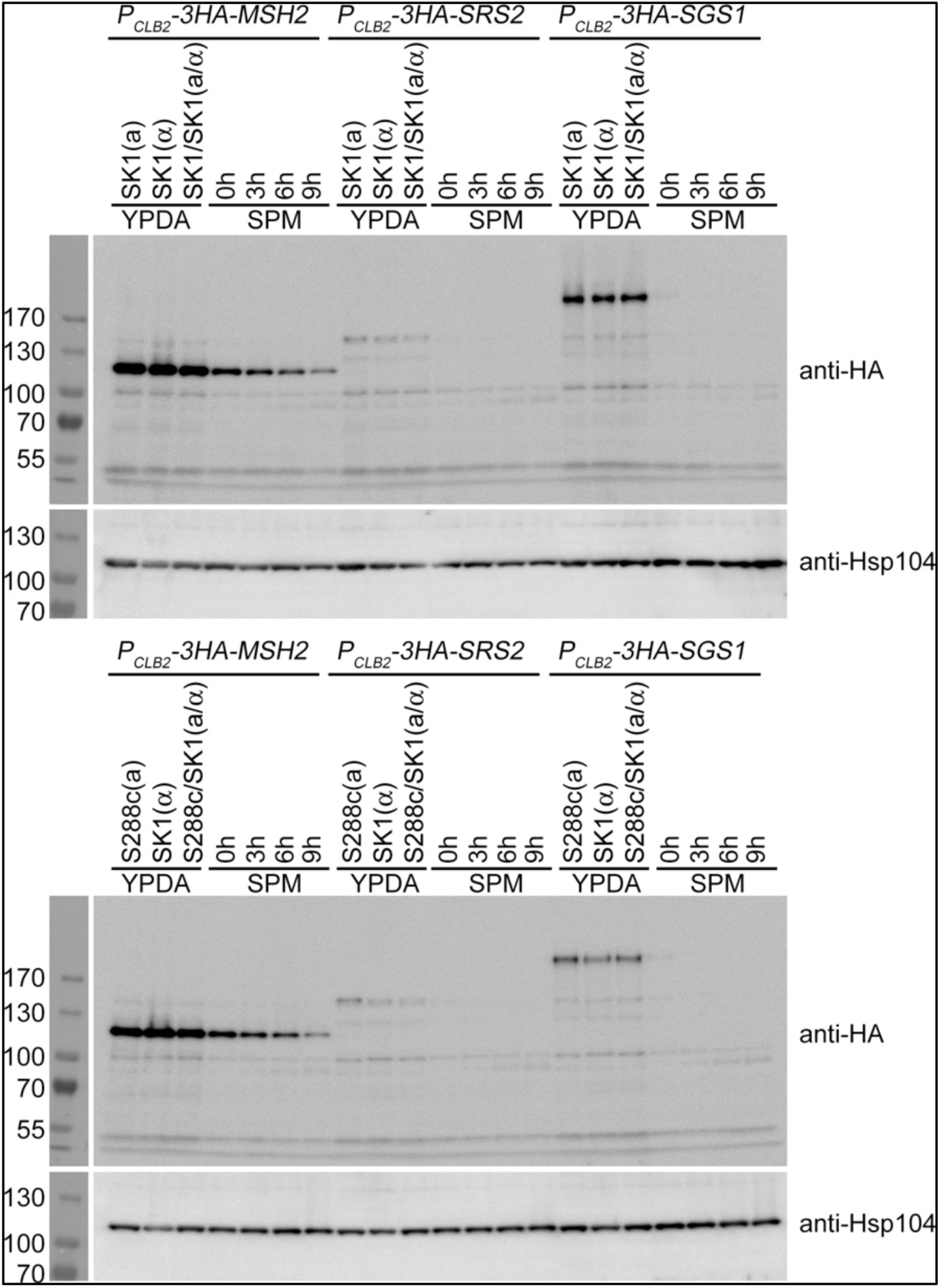
*P_CLB2_* represses meiotic expression of *MSH2*, *SGS1* and *SRS2*. Whole-cell extracts of ∼6 × 10^7^ vegetative or sporulating cells were resuspended with 1 ml of freshly prepared buffer (0.185 M NaOH, 0.75% β-mercaptoethanol), incubated on ice for 15 min, and mixed with 150 µl of 55% trichloroacetic acid (TCA) for 10 min on ice. The cell suspension was centrifuged at 13,500 rpm for 10 min. The protein pellets were resuspended with HU buffer (8 M urea, 5% SDS, 200 mM Tris-HCl pH 6.5, 5 mM EDTA, 0.01% bromophenol blue and 4.5% β-mercaptoethanol). Protein samples were denatured at 65 °C for 10 min and then separated by SDS-PAGE. Vegetative haploid and diploid cells were grown at 30 °C in YPDA medium to OD_600nm_ = 0.4 -0.6. Sporulating cells were first grown at 30 °C overnight in YPA medium (1% yeast extract, 2% peptone, and 2% potassium acetate or KAc), washed twice with sterilized double-distilled water (ddH_2_O), transferred into sporulation medium (2% KAc) at the zero time-point, and then harvested at the indicated time-points. Supplementary Dataset DS21 contains uncropped blots of Figure S1.

**Figure S2.**
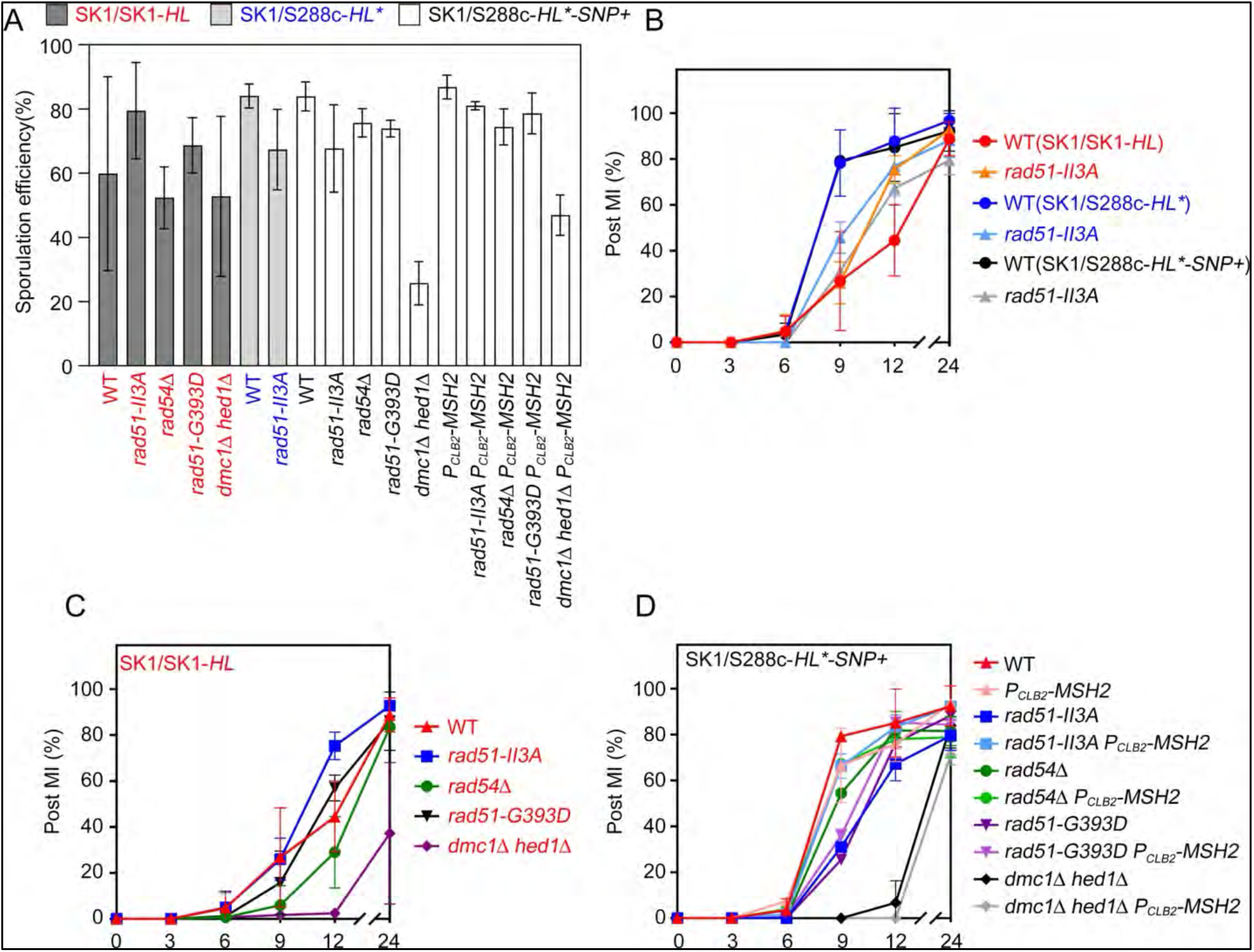
Sporulation efficiency (A) and meiotic progression (B-D) of the hybrid zygotes analyzed in Figure 6E.

**Figure S3.**
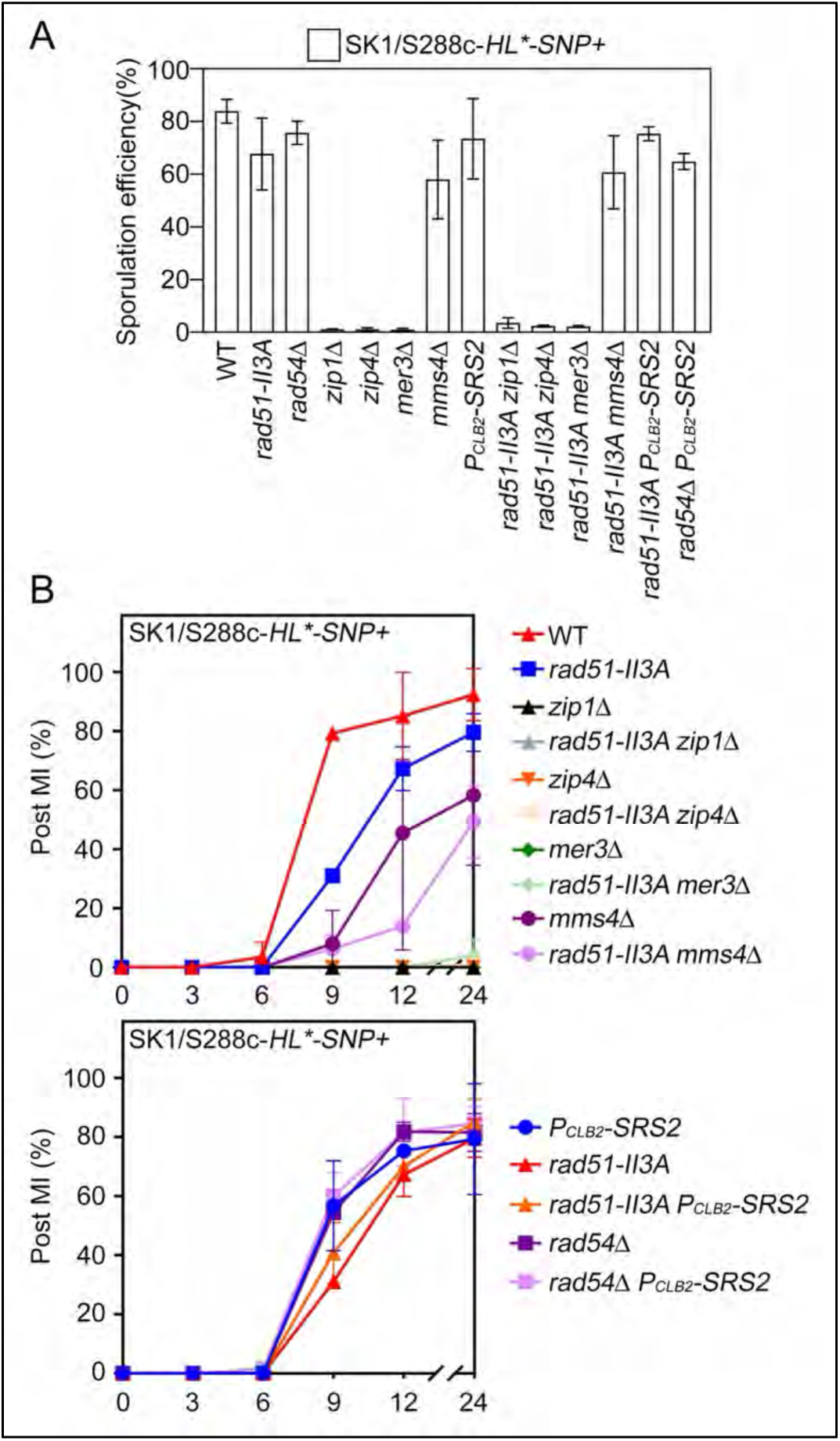
Sporulation efficiency (A) and meiotic progression (B) of the hybrid zygotes analyzed in Figure 6F.

**Figure S4.**
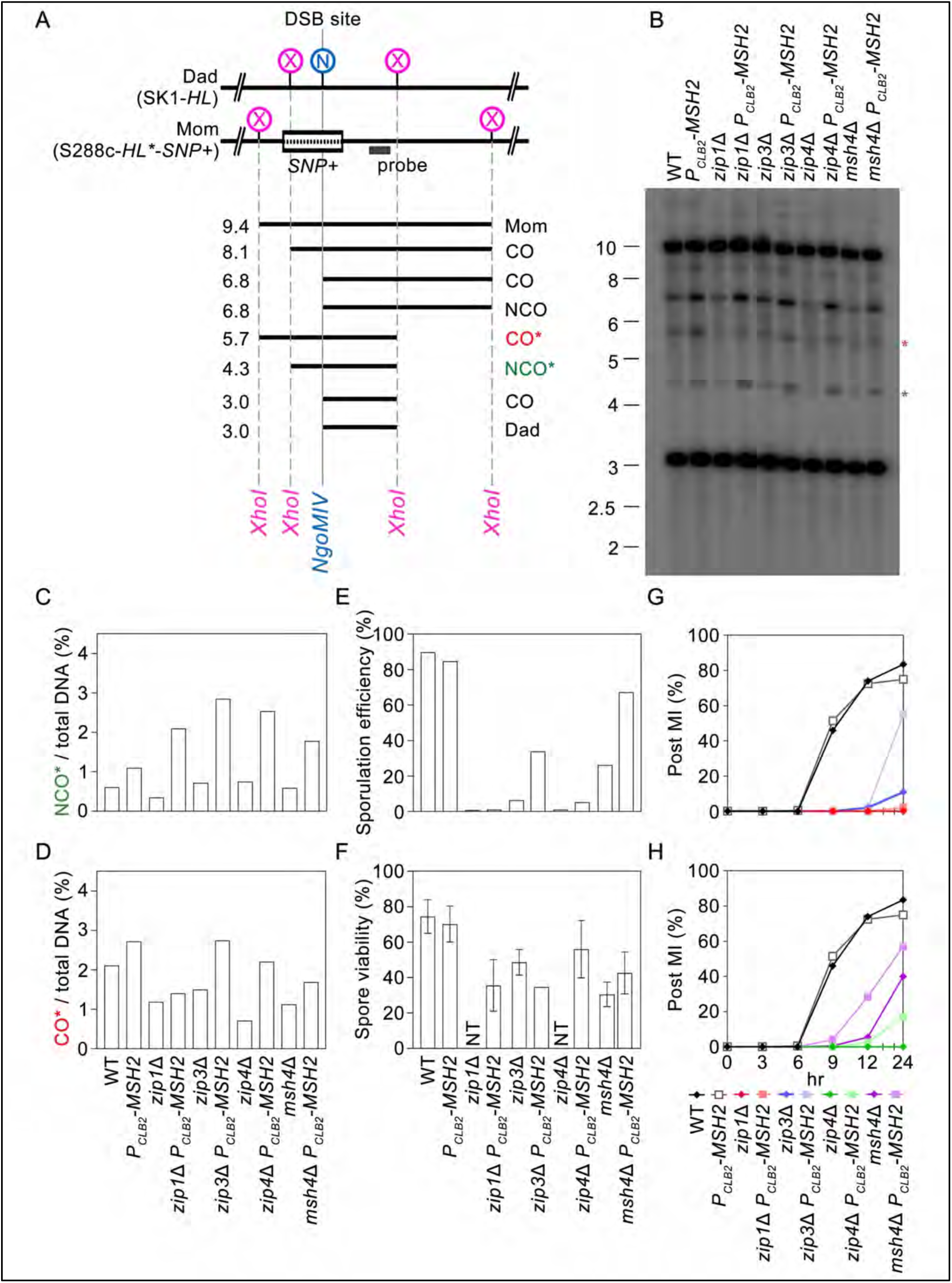
The effect of *P_CLB2_-MSH2* on the *zmm*/1 SK1/S288c-*HL*\**-SNP+* hybrid, including the quantities of recombinants at the *HL*\**-SNP+* DSB hotspot (A-D), sporulation efficienty (E), spore viability (F), and progression of MI (G-H)).

**Figure S5.**
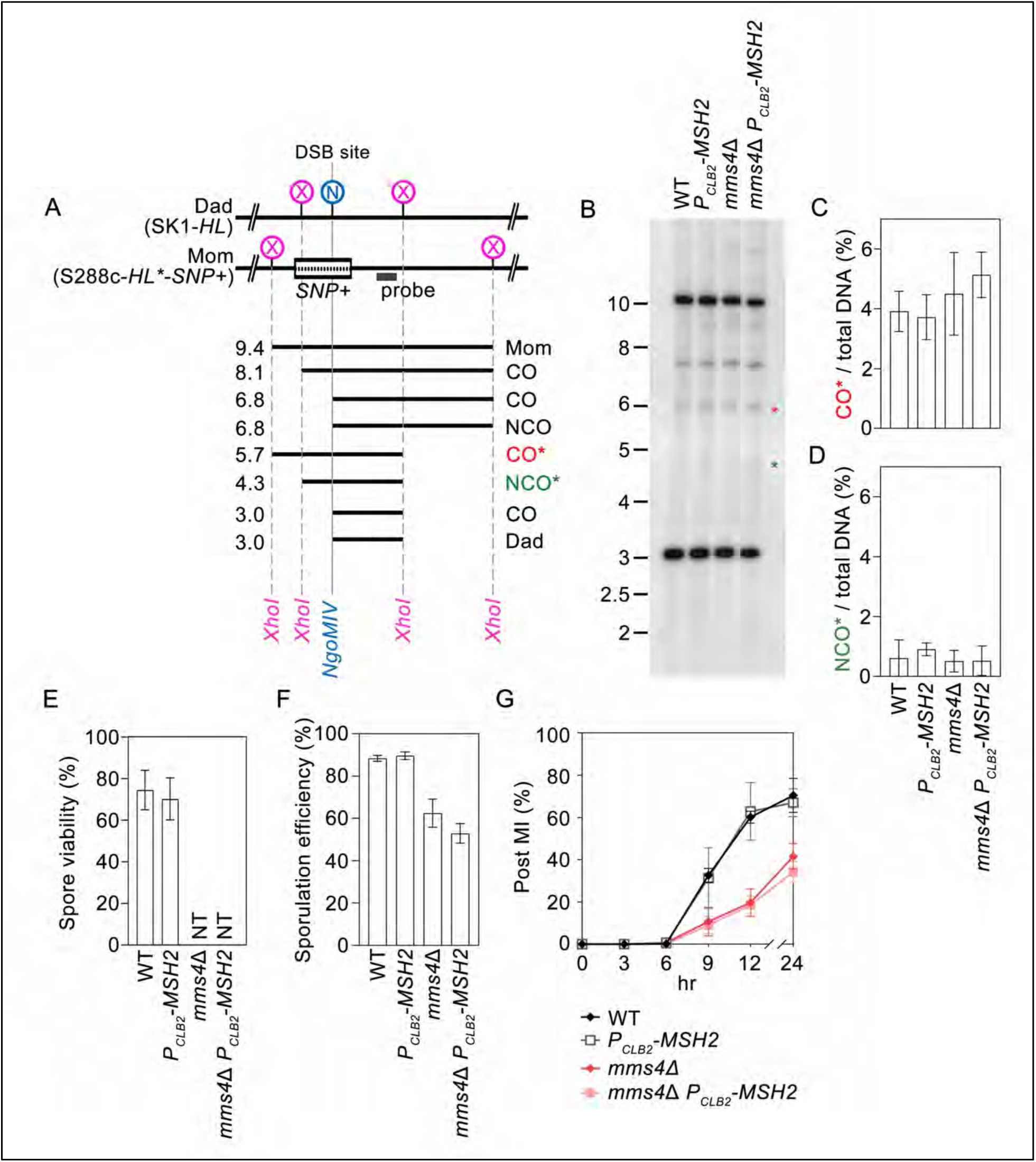
The effect of *P_CLB2_-MSH2* on the *mms4*/1 SK1/S288c-*HL*\**-SNP+* hybrid, including the quantities of recombinants at the *HL*\**-SNP+* DSB hotspot (A-D), spore viability €, sporulation efficiency (F), and meiotic progression (G).

**Figure S6.**
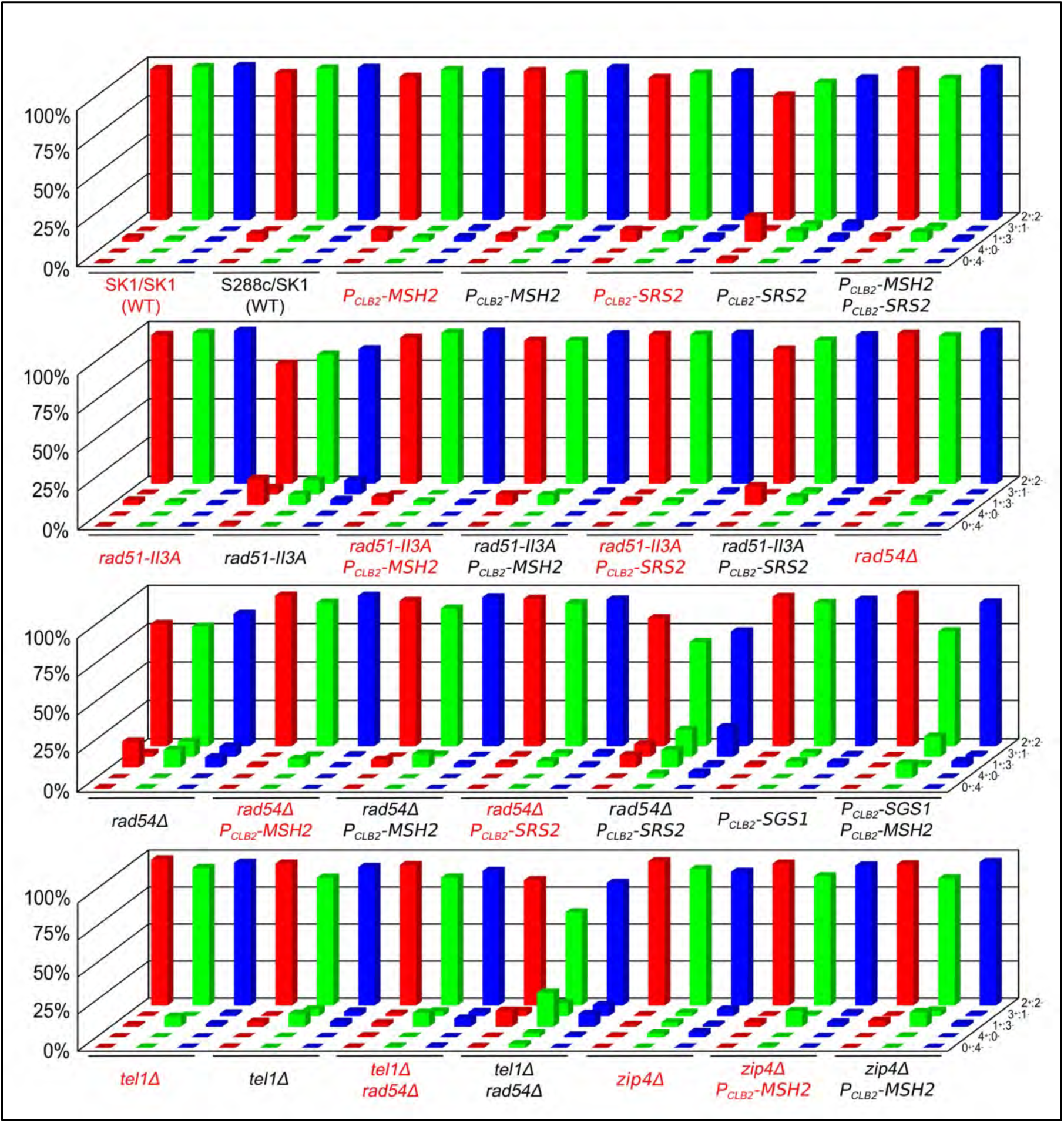
The percentages of tetrads underwent 2^+^:2^−^ 3^+^:1^−^, 1^+^:3^−^, 4^+^:0^−^, or 0^+^:4^−^. segregation from each pair of heterozygous spore-autonomous fluorescent marker genes. *CEN8::RFP/CEN8* (in red); *ARG4::GFP*-*ARG4* (in green); *THR1::CFP*-*THR1* (in blue).

**Table S1.** List of DNA repair and recombination mutants analyzed in this study.

| # | Mutant | Description |
| --- | --- | --- |
| 1 | *dmc1Δ* | A meiosis-specific RecA-like recombinase (1). |
| 2 | *rad51Δ* | A ubiquitous RecA-like recombinase (2). |
| 3 | *rad51-ΔN* | An extremely labile mutant lacking the N-terminal domain (NTD, 1-66 a.a.). Removal of NTD from Rad51 (*Rad51-ΔN*) reduces by >95% the steady-state protein levels, but the *Rad51-ΔN* mutant retains most of its DSB repair activities during vegetative growth and meiosis (3). |
| 4 | *rad51-NTD3A* | A phosphorylation-defective mutant with a shorter half-life (3). |
| 5 | *dmc1Δ hed1Δ* | Hed1 is a meiosis-specific inhibitor that acts by preventing Rad51 from interacting with Rad54. This mutant possesses a catalytically active Rad51 (4-7). |
| 6 | *dmc1-P267V* | The *Dmc1-P267V/G268D* mutant lacks *Dmc1*'s capability in tolerating DNA mismatch during homology pairing and the strand exchange reaction (8). We show in this study that this mutant behaves similarly to wild-type *Dmc1* in both SK1/SK1 and SK1/S288c zygotes (data not shown). |
| 7 | *dmc1-G268D* | The *Dmc1-P267V/G268D* mutant lacks *Dmc1*'s capability in tolerating DNA mismatch during homology pairing and the strand exchange reaction (8). We show in this study that this mutant behaves identically to *dmc1Δ* in both SK1/SK1 and SK1/S288c zygotes (Table S3). |
| 8 | *dmc1-P267V,G268D* | The *Dmc1-P267V/G268D* mutant lacks *Dmc1*'s capability in tolerating DNA mismatch during homology pairing and the strand exchange reaction (8). We show in this study that this mutant behaves identically to *dmc1Δ* in both SK1/SK1 and SK1/S288c zygotes (Table S3). |
| 9 | *rad51-V331P*<br>*dmc1Δ hed1Δ* | The *Rad51-V331P/D332G* mutant has been reported to behave like *Dmc1* in tolerating DNA mismatch during homology pairing and the strand exchange reaction (8). However, we show in this study that *rad51-V331P* cannot rescue the low spore viability phenotype of SK1/S288c *dmc1Δ hed1Δ* hybrid zygotes (Table S3). |
| 10 | *rad51-D332G*<br>*dmc1Δ hed1Δ* | The *Rad51-V331P/D332G* mutant has been reported to behave like *Dmc1* in tolerating DNA mismatch during homology pairing and the strand exchange reaction (8). However, we have shown that *rad51-D332G* cannot rescue the low spore viability phenotype of SK1/S288c *dmc1Δ hed1Δ* hybrid zygotes (Table S3). |
| 11 | *rad51-V331P, D332G*<br>*dmc1Δ hed1Δ* | The *Rad51-V331P/D332G* mutant has been reported to behave like *Dmc1* in tolerating DNA mismatch during homology pairing and the strand exchange reaction (8). However, we have shown that *rad51-V331P/D332G* cannot rescue the low spore viability phenotype of SK1/S288c *dmc1Δ hed1Δ* hybrid zygotes (Table S3). |
| 12 | *rad51-II3A*<br>(*R188A/K361A/K371A*) | A *Rad51* mutant with a defective DNA binding site II, lacking homology pairing and strand exchange activity (9). |
| 13 | *rad52Δ* | *Rad52* is involved in both *Rad51*-dependent and *Rad51*-independent homologous recombination. *Rad52* attenuates the resection of DNA DSB ends and stimulates strand exchange by facilitating binding of phosphorylated *Rad51* (*Rad51p*) to single-stranded DNA (ssDNA). *Rad52* anneals complementary ssDNA. *Rad52* protects *Rad51* nucleoprotein filaments against anti-recombinogenic *Srs2* DNA helicase (10-12). |
| 14 | *rad54Δ* | A DNA-dependent ATPase that stimulates Rad51-mediated strand exchange (13,14). Rad54 protects Rad51 nucleoprotein filaments against anti-recombinogenic Srs2 DNA helicase (12). |
| 15 | *rdh54Δ (tid1Δ)* | A DNA-dependent ATPase that stimulates Dmc1-mediated strand exchange (15-18). |
| 16 | *rad51-A320V* | A Rad51 mutant defective in the Rad51-Rad52 interaction. It retains the ability (equivalent to wild-type) to interact with Rad54, Rdh54, and Rad59 (10). |
| 17 | *rad51-Y388H* | A Rad51 mutant defective in the Rad51-Rad52 and Rad51-Srs2 interactions. It retains the ability (equivalent to wild-type) to interact with Rad54, Rdh54, and Rad59. The Rad51-Y388H-ssDNA presynaptic filament is resistant to Srs2 (10). |
| 18 | *rad51-G393D* | A Rad51 mutant defective in the Rad51-Rad52 and Rad51-Srs2 interactions. It retains the ability (equivalent to wild-type) to interact with Rad54, Rdh54, and Rad59. The Rad51-G393D-ssDNA presynaptic filament is resistant to Srs2 *in vitro* (10). |
| 19 | *dmc1-A256V* | The parallel mutant of *rad51-A320V*. |
| 20 | *dmc1-G325D* | The parallel mutant of *rad51-Y388H*. |
| 21 | *dmc1-T330D* | The parallel mutant of *rad51-G393D*. |
| 22 | *rad51-E221D* | The parallel mutant of *dmc1-E157D*. |
| 23 | *rad51-D239D* | A Rad51 mutant defective in binding to the Rad55•Rad57 complex (19) (Table S3). |
| 24 | *rad51-D241A* | A Rad51 mutant defective in binding to the Rad55•Rad57 complex (19) (Table S3). |
| 25 | *dmc1-E173A* | The parallel mutant of *rad51-D239A*. |
| 26 | *dmc1-D175A* | The parallel mutant of *rad51-D239A*. |
| 27 | *dmc1-D177A* | The parallel mutant of *rad51-D241A*. |
| 28 | *dmc1-E157D* | Dmc1-E147D is uncoupled from the Mei5•Sae3 complex (20). |
| 29 | *rad59Δ* | Rad59 is a paralog of Rad52 that is required for Rad51-independent HR. It promotes the loading of phosphorylated Rad52 (Rad52p) at DSBs and initiates the single strand annealing (SSA) DSB repair pathway (21). |
| 30 | *ku70Δ* | Ku70 and Ku80 form a heterodimer involved in the canonical nonhomologous end joining (NHEJ) DSB repair pathway (22). |
| 31 | *msh2Δ* | In eukaryotes, mismatch recognition is accomplished by two heterodimeric MutS homolog complexes, Msh2•Msh3 (MutSα) and Msh2•Msh6 (MutSβ) (23-25). |
| 32 | *P<sub>CLB2</sub>-MSH2* | *P<sub>CLB2</sub>* is the native promoter of the mitosis-specific B-type cyclin *CLB2*, which is specifically repressed during meiosis (26). |
| 33 | *P<sub>CLB2</sub>-MSH3* | Msh2 and Msh3 form a heterodimeric MutSα homolog complex that acts primarily on insertion/deletion loop mismatches up to 12 nucleotides in length (23-25). |
| 34 | *P<sub>CLB2</sub>-MSH6* | Msh2 and Msh6 form a heterodimeric MutSβ homolog complex that is primarily involved in repairing base-base and small insertion/deletion loop mismatches (23-25). |
| 35 | *pms1Δ* | Pms1 and Mlh1 form a heterodimeric MutSα homolog complex (27). |
| 36 | *mlh2Δ* | Mlh2 and Mlh1 form a heterodimeric MutLβ homolog complex (27). |
| 37 | *msh3Δ* | Mlh3 and Mlh1 form a heterodimeric MutLγ homolog complex (27). |
| 38 | *P<sub>CLB2</sub>-SRS2* | Srs2, the *S. cerevisiae* homolog of *E. coli* UvrD (also called MutU), binds to Rad51 (not Dmc1) and disassembles the Rad51-ssDNA presynaptic filament (28-30). Srs2 physically interacts with Mms4•Mus81 and promotes Mms4•Mus81-mediated resolution of recombination intermediates (31). Srs2's ATPase and ssDNA translocating motor activities are inhibited by Dmc1 (32). The hyper-gene conversion Hpr5-1 mutation is an allele of the *SRS2* gene (33). |
| 39 | *P<sub>CLB2</sub>-MPH1* | Mph1, the *S. cerevisiae* ortholog of mammalian Fanconi anemia helicase, prevents precocious DSB strand exchange between sister chromatids before homologs have completed pairing, thereby ensuring high levels of IH-COs and IH-NCOs between homologs (34). |
| 40 | *P<sub>CLB2</sub>-PIF1* | Pif1 specifically facilitates Rad51-mediated IH-HR but not Dmc1-mediated IH-HR (35). Pif1 also stimulates DNA polymerase delta (Pol δ)-mediated DNA synthesis from D-loops through its interaction with proliferating cell nuclear antigen (PCNA), a sliding clamp that increases the processivity of all DNA polymerases (36). |
| 41 | *P<sub>CLB2</sub>-MMS4*<br>*mms4Δ* | The Mms4•Mus81 endonuclease complex collaborates with Sgs1 to direct meiotic recombination toward interhomolog interactions that promote proper chromosome segregation. Mms4•Mus81 promotes joint molecule (JM) resolution *in vivo* (37,38). |
| 42 | *sgs1Δ* | *S. cerevisiae* has one RecQ helicase, Sgs1, which functions with Top3 and Rmi1 in DNA repair. Mutations in the *SGS1* gene lead to defects similar to those seen in human cells displaying RecQ family disorders, such as increases in the frequency of recombination and chromosome loss (39,40). The *sgs1Δ* null mutation increases the rate of chromosome synapsis, displays an elevated level of crossing over, and triggers checkpoint-induced arrest at pachytene (41). |
| 43 | *sgs1ΔC795* | The *sgs1-ΔC795* allele removes the carboxy-terminal 795 amino acids. The *sgs1-ΔC795* mutant grows as well as wild-type and displays a level of hydroxyurea sensitivity intermediate between wild-type and *sgs1Δ* but sporulates more efficiently than the null mutants and its vegetative growth rate is essentially normal (41). |
| 44 | *P<sub>CLB2</sub>-SGS1* | Early NCO formation is abolished in *S. cerevisiae* mutants lacking Sgs1, the ortholog of mammalian Bloom RecQ-like helicase (42). Sgs1 also acts downstream of mismatch recognition by Msh2 to unwind nascent recombination intermediates containing a high density of heteroduplex basepair mismatches (HDBMs) during multiple DNA repair steps (43). |
| 45 | *zip1Δ* | Zip1, a ZMM family protein, is the transverse filament protein of the synaptonemal complex. Zip1 is required for normal levels of meiotic recombination and pairing between homologous chromosomes during meiosis (44). Zip1 also mediates homology-independent centromere coupling during early meiosis (45). |
| 46 | *mer3Δ* | Mer3, a ZMM family protein, is a DNA helicase with two meiotic functions. It competes with Pif1 for binding to the PCNA clamp loader RFC, thereby limiting polymerase delta (Pol δ)-mediated DNA synthesis during DSB repair (46). It also interacts with Dmc1 to antagonize the anti-recombinase activities of Sgs1 (47). |
| 47 | *zip4Δ* | Zip4 (Spo22) forms a heterotrimeric ZZS complex with Zip2 and Spo16. |
|  |  | The ZZS complex exerts two distinct activities: Zip4 acts as a hub for physical interactions with components of the chromosome axis and the crossover machinery, whereas Zip2 and Spo16 have non-catalytic DNA recognition functions essential to CO formation (48,49). |
| 48 | *zip3Δ* | Zip3 is a small ubiquitin-like modifier (SUMO) E3 ligase that marks the sites of COs (50-53). |
| 49 | *msh4Δ* | Msh4 and Msh5 are two meiosis-specific MutS proteins that form the MutSγ heterodimeric complex. MutSγ is not involved in mismatch recognition, but facilitates crossing over by binding and stabilizing nascent recombination intermediates (49,54,55). |
| 50 | *ecm11Δ* | A meiosis-specific component of the synaptonemal complex along with Gmc2. The Ecm11•Gmc2 complex is required for efficient crossover formation and for SC assembly (56). |
| 51 | *zip4-N919Q* | Zip4 brings Ecm11 to recombination sites that are prone to form COs and helps the transverse filament protein Zip1 to nucleate from this location, ensuring control of recombination initiates locally from sites engaged in the crossover repair pathway. The N919Q point mutant specifically abolishes the Zip4-Ecm11 interaction, leading to loss of Ecm11 binding to chromosomes and defective SC assembly (57). |
| 52 | *zip2-ΔXPF* | Spo16 binds the C-terminal XPF domain of Zip2 to form a meiosis-specific complex that shares structural similarities to the structure-specific XPF–ERCC1 nuclease, although it lacks endonuclease activity. The *zip2-ΔXPF* mutant encodes a Zip2 mutant protein lacking the XPF domain. Despite a minor difference between CO levels, sporulation efficiency and spore viability of the *zip2-ΔXPF* mutant are significantly higher than for the *zip2Δ* mutant, suggesting that the N-terminal part of Zip2 also participates in its pro-CO activity (48). |

**Table S2.**
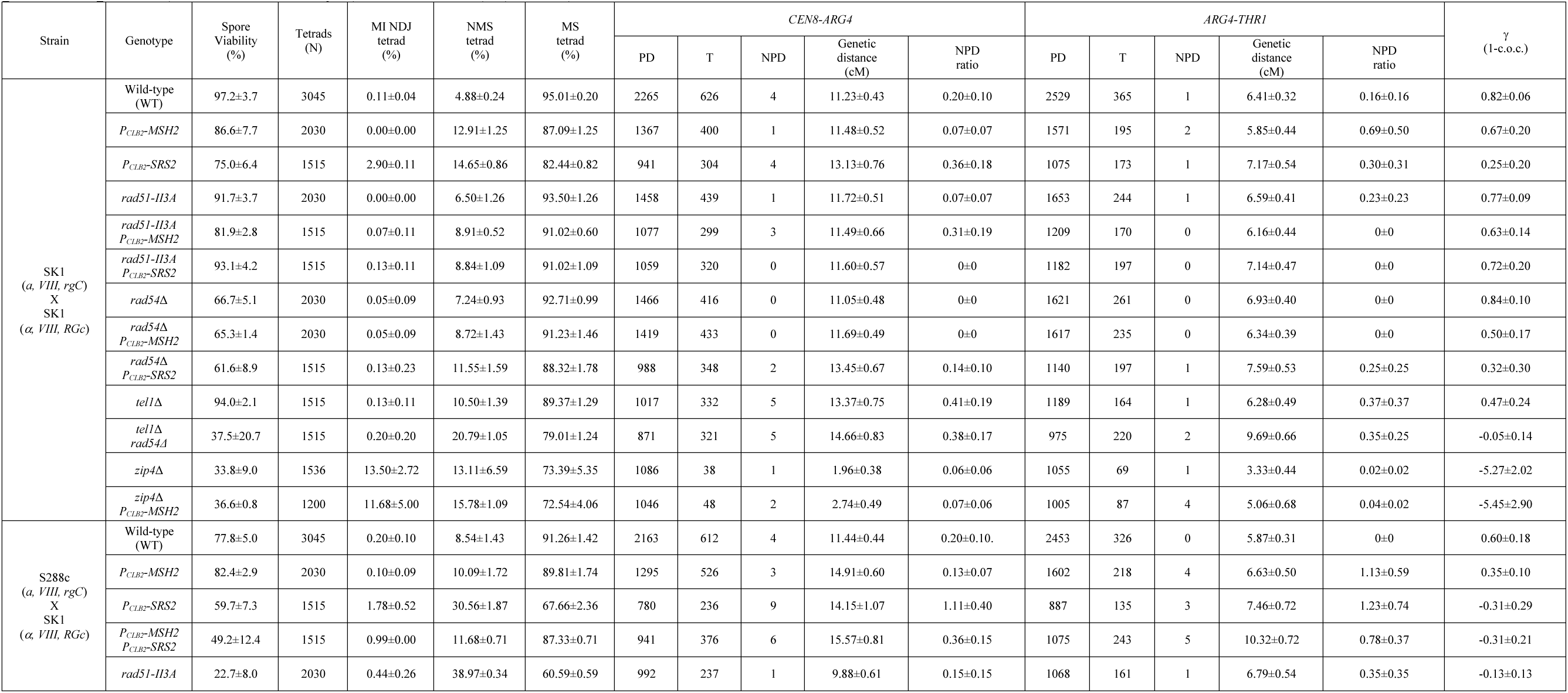
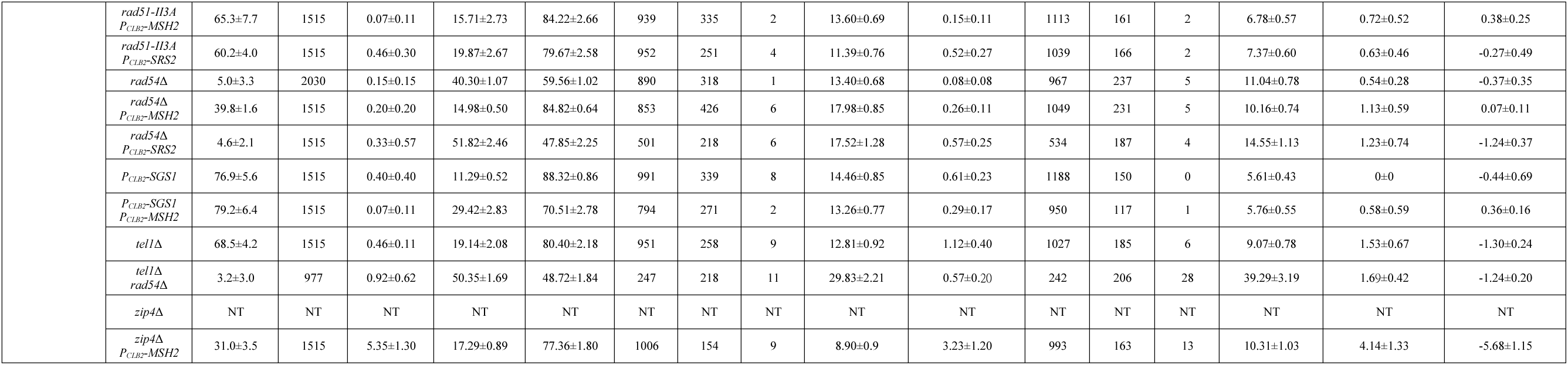
Genetic distances and CO interferences. Intraspecies SK1(*a, VIII, rgC*)/SK1(*α, VIII, RGc*) purebred zygotes and intraspecies S288c(*a, VIII, rgC*)/SK1(*α, VIII, RGc*) hybrid zygotes host three heterozygous marker genes [*CEN8*(*r*)*/CEN8::RFP*(*R*)*, ARG4*(*g*)*/ARG4::GFP*(*G*), and *THR1::CFP*(*C*)*/THR1*(*c*)] on chromosome VIII (138). The diploid zygotes were used to generate tetrads and determine spore viability, the percentages of tetrads with MI NDJ, NMS (with ≥1 non-2^+^:2^−^ fluorescent protein markers) and MS (with three 2^+^:2^−^ fluorescent protein markers), respectively. Higher NMS percentages indicate higher IH-NCO or allelic GC rates. The MS tetrads were used to calculate CO interference (γ), genetic distances and the “NPD ratio” in two genetic intervals using classical genetic methods (https://en.wikipedia.org/wiki/Coefficient_of_coincidence) (74) and Stahl lab’s online tool (https://elizabethhousworth.com/StahlLabOnlineTools/). A positive interference value indicates spacing of IH-CO events that departs from a random distribution, whereas negative interference describes IH-CO events that are more clustered than the null expectation. The “NPD ratio” (fN_obs_/fN_exp_) is an indication of the intensity of interference, with smaller values indicating stronger positive interference. *P_CLB2_* is the native promoter of the mitosis-specific B-type cyclin gene *CLB2* that is deactivated during meiosis. N: the total number of scored tetrads. NT: not enough mature tetrads. We combined three independent sets of each strain and scored tetrads either individually (N/3; split in triplicate) or collectively (i.e., N; total) (DS23).

